# Using a planted tree biodiversity experiment to evaluate imaging spectroscopy for species classification

**DOI:** 10.64898/2026.03.20.713086

**Authors:** Sofia J. van Moorsel, Bernhard Schmid, Michael Niederberger, Jeremiah Huggel, Michael Scherer-Lorenzen, Uwe Rascher, Alexander Damm, Meredith C. Schuman

## Abstract

Field-based monitoring of tree species in forests is often sparse due to logistical constraints. Remote sensing enables repeated, spatially contiguous collection of reflectance data across large areas. Tree species classification accuracy using such data is variable, likely because most studies use observational datasets where species occurrence correlates with environmental variation. We used two sites of a tree biodiversity experiment in Germany (BIOTREE: Kaltenborn and Bechstedt), where different species have been planted with high replication under controlled diversity levels, to assess how well tree species could be classified using reflectance data from airborne imaging spectroscopy and different classification methods (linear discriminant analysis, LDA, and a non-linear support vector machine, SVM). Reflectance data for 589 wavelengths between 400–2400 nm were acquired at 1 m spatial resolution during peak growing season. Reflectance spectra showed large and significant variation between taxonomic classes, orders, and species, and weak, but still significant, interactions between classes or orders and diversity levels. Classification accuracy reached 100 % in training datasets, 77 %-83 % for the four species in Kaltenborn prediction datasets, and 31 %-49 % for the 16 species in Bechstedt prediction datasets. LDA provided more accurate predictions than SVM; and using similarly-spaced original wavelengths with LDA was as efficient as using principal components derived from the original data. While airborne imaging spectroscopy effectively distinguished up to four tree species in our datasets, classification accuracy was lower when 16 species had to be distinguished. In these cases, the methodology may be more useful for functional diversity monitoring than for tree species classification.

## 1. Introduction

Forests cover approximately one third of Earth’s land surface and harbor 80 % of global terrestrial biodiversity (FAO and UNEP, 2020). Forest biodiversity maintains ecosystem health and resilience; it is the basis of forest productivity and ensures the provision of ecosystem goods and services vital to humanity (Liu et al., 2026). However, forest biodiversity is in decline (IPBES, 2019). We require efficient methods for the monitoring of forest biodiversity across large, contiguous areas to understand dynamics of biodiversity loss and facilitate management and protection. Spectroscopy, a sub-field of remote sensing, offers a scalable solution to map forest functional diversity (Hauser et al., 2021; Aguirre-Gutiérrez et al., 2021; Schneider et al., 2017) and – by enabling species identification (Klehr et al., 2025; Hermosilla et al., 2024) – tree species richness. However, matching sensor type to ecological scale is crucial: leaf probes capture individual traits, while aircraft-based measurements are well suited for local community-scale studies, and satellites provide repeated coverage at regional to global scales (Cavender-Bares et al., 2025a).

Imaging spectroscopy has long been recognized as a powerful approach for tree species discrimination and biodiversity assessment because reflectance spectra capture variation in leaf chemistry, canopy structure, water content, and physiology that is often taxonomically structured (Asner and Martin, 2009; Cavender-Bares et al., 2025b). Early studies demonstrated species separability using airborne hyperspectral sensors such as AVIRIS in both temperate and tropical forests (Gong et al., 1997; Townsend et al., 2003; Clark et al., 2005). Subsequent work expanded these approaches to species mapping, functional trait estimation, and biodiversity assessment across diverse ecosystems using imaging spectroscopy and combinations of hyperspectral and LiDAR data (Baldeck et al., 2015; Somers and Asner, 2013; Roth et al., 2015; Seeley et al., 2024; Tagliabue et al., 2020). Despite these advances, most of these studies have relied on sample surveys rather than experiments (Liu et al., 2026). In sample surveys, species occurrence co-varies with environmental gradients, canopy position, stand structure, and neighborhood composition, making it difficult to disentangle species-specific spectral signatures from environmental variation. For example, if species A tends to grow on ridges and species B in valleys, then the remote sensing signal may actually discriminate between ridges and valleys rather than between species A and B, or at least its ability to distinguish the species may be contingent on their environments.

Tree species classification often relies on high-resolution (cm-m) true color (RGB) or multispectral images, for example, to derive phenological information (*e.g.*, green-up and senescence) during the course of multiple acquisitions over the growing season (Fawcett et al., 2021; Hardenbol et al., 2021). If sensors are installed on uncrewed airial vehicles (UAVs) or piloted airplanes, hyperspectral imaging spectroscopy is typically restricted to smaller spatial extents. Satellite data offer broader spatial coverage, but may lack the spatial and spectral resolution needed to distinguish species in mixed forests (but see Koch et al. (2025)). Sensors mounted on airplanes provide an intermediate solution, covering larger areas than UAVs, yet offering higher spatial resolution than public (and most private) satellites for the same or better spectral resolution (Chadwick et al., 2025; Schaepman et al., 2015; Hueni et al., 2025). Airborne campaigns have been used to map invasive species, assess ecosystem recovery, and estimate functional diversity (Schneider et al., 2017, 2023; Vögtli et al., 2025). Airplanes also offer potential for interdisciplinary ecosystem research since they can host multiple instruments and multi-investigator deployments of longer duration (Morsdorf et al., 2020). However, due to the relatively high cost of airborne campaigns, it is critical to evaluate whether a single-acquisition airborne survey can provide meaningful data on species identities and thus species richness at the community level in forests.

A particular challenge for tree species identification using imaging spectroscopy in natural forests is that different species may have similar spectral properties (e.g., reflectance spectra) due to shared traits, while members of the same species may have different reflectance spectra because they grow on different microsites or with different neighbors, which can mask the differences they would show when planted in the same environment and with the same neighbor species (Liu et al., 2026). The impact of such spectral confusion within and across species, and thus the potential of imaging spectroscopy to differentiate species, can be best assessed in forests planted across shared environmental conditions and with designed species arrangements, as is typically done in forest biodiversity experiments.

Here, we use two sites of the German BIOTREE experiment (Scherer-Lorenzen et al., 2007) to assess the discriminatory power of reflectance spectra obtained from single airplane overflights for tree species classification. The advantage of this particular experiment is that species have been planted in small monospecific patches, assembled into forest stands consisting of different numbers of species or different functional diversity. Like this, we could obtain top-of-canopy reflectance spectra at pixel level over known species without needing to perform crown delineation.

We first tested how the reflectance spectra of the tree patches were affected by environmental (different plots and different tree diversities) and species differences with their interactions using general linear models across 589 spectral bands in the 400–2500 nm wavelength region. In this analysis, we identified spectral ranges that are most discriminatory regarding environmental and species differences. We then used linear discriminant analysis (LDA) and a non-linear machine-learning approach (a support-vector machine, SVM) implemented in R (Meyer et al., 2024) to classify the tree species. We used different numbers of similarly-spaced wavelengths or different numbers of principal components derived from all wavelengths by principal component analysis (PCA). We compare classification accuracy in training datasets and prediction datasets for the different methods used at the two experimental sites, Kaltenborn with four and Bechstedt with 16 tree species. We offer the example dataset as a reference for further method tests and comparisons.

## 2. Material and Methods

### 2.1. Biodiversity experiment sites

This research was conducted on sites of the BIOTREE experiment established in 2002 in Thuringia, Germany (Fig. 1). For detailed information on the site characteristics, see (Scherer-Lorenzen et al., 2007). In both experiments trees were planted in monoculture patches and these patches combined to diversity treatments at plot level. In Kaltenborn, the patches were mixed to create different levels of species richness (1, 2, 3, and 4). In Bechstedt, the richness level of the plots was always 4 species. However, the four species were selected in such a way that they were functionally more or less similar, yielding a range of functional diversity levels while holding richness constant at 4 species. To create different functional diversities at plot level, we first classified the 16 species used as species pool into different functional groups using a cluster analysis with nine above- and belowground functional traits that represented attributes indicative for productivity, resource use, and nutrient cycling (e.g., leaf type, light requirement, mean annual increment, leaf N concentration, litter C/N ratio). We then assembled different sets of four species (in total 1.820 possibilities) such that they represented a range of functional richness from highly similar to highly different (24 different sets used in the experiment, with one species combination planted twice). We quantified the functional diversity using a dendrogram-based index FD (Petchey and Gaston, 2002).

**Fig. 1.**
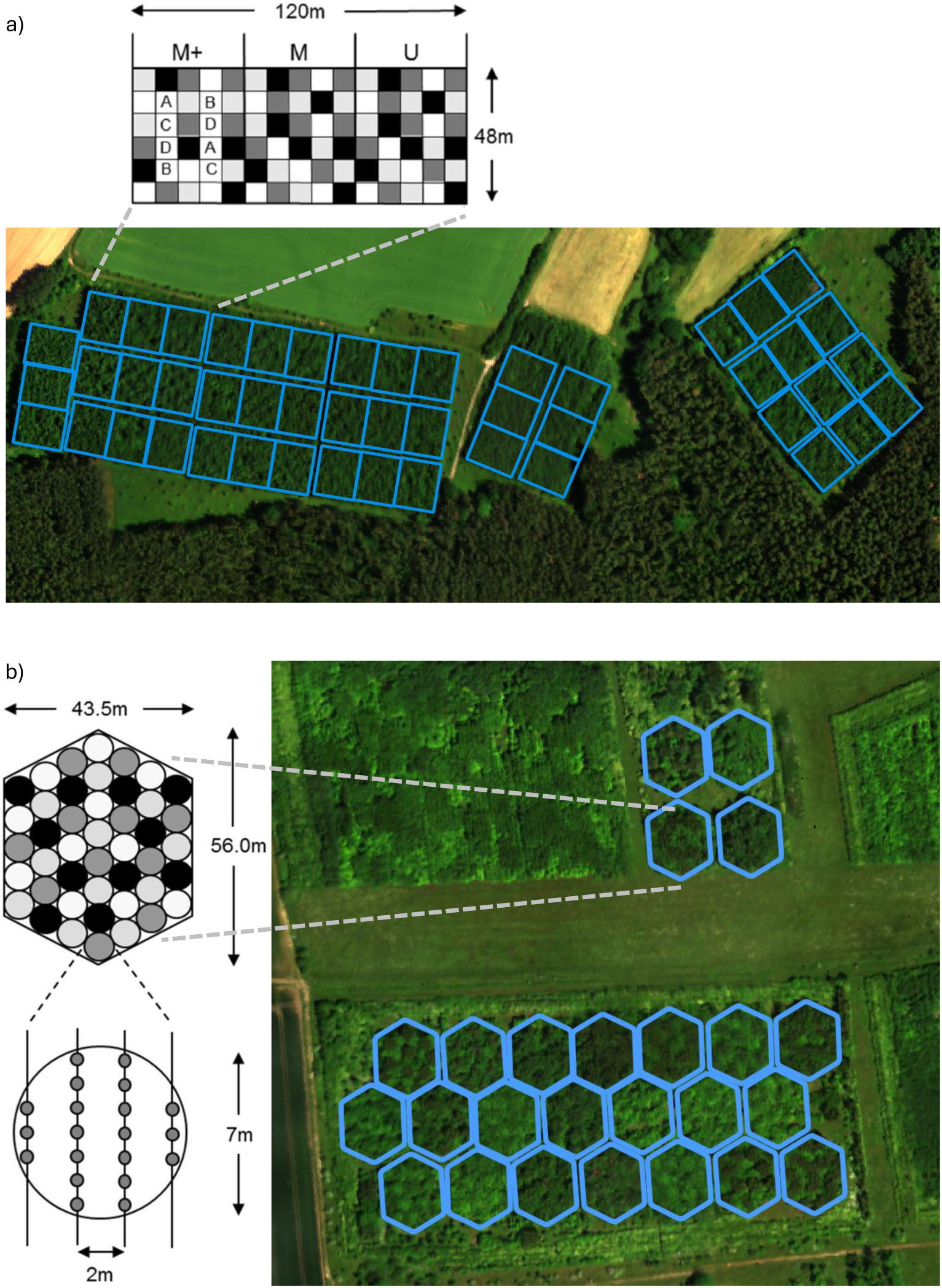
Overview of investigated test sites (true colour HyPlant images). a) Kaltenborn field site with 16 rectangular plots and the planting design for one of these. Trees of a single species were planted together in quadratic patches of 8 x 8 m. b) Bechstedt field site with 25 hexagonal plots and the planting design shown for one of these. Within each plot, the species were planted in circular plots of 3.5 m radius containing 20 individuals of the same species.

#### 2.1.1. Kaltenborn

The “BIOTREE-SPECIES: Kaltenborn” site (50.77867, 10.22259, WGS84) contains 16 plots of 120 x 48 m with different species compositions (Fig. 1a), each split into three subplots of 40 x 48 m with different forest management strategies: unmanaged (u), managed (thinned according to regional state-of-the-art management rules of close-to-nature forestry, labeled m), and managed with the addition of four species at lower abundance than the main species (m+, (Scherer-Lorenzen et al., 2007)). In this study, we included the treatments unmanaged (u) and managed (m). Trees of a single species were planted in 8 x 8 m patches (30 patches per subplot for a total of 960 patches included in the analyses) which were then randomly combined in a checkerboard manner to obtain the designed species composition with one, two, three, or four species per plot. The 16 different species compositions were assembled from a pool of four common tree species: the two gymnosperms Douglas fir (*Pseudotsuga menziesii*, n = 242 patches) and Norway spruce (*Picea abies*, n = 242 patches, two outliers excluded for further analysis) and the two angiosperms European beech (*Fagus sylvatica*, n = 238 patches) and sessile oak (*Quercus petraea*, n = 238 patches).

#### 2.1.2. Bechstedt

At the “BIOTREE-SPECIES: Bechstedt” site (50.8971, 11.08238, WGS84), only functional diversity was manipulated, and the site contains 16 different species (Scherer-Lorenzen et al., 2007) in a total of 25 hexagonal plots (Fig. 1 b). Within each plot, four species were planted in monospecific circular patches of 3.5 m radius containing 20 individuals in four rows (between-row distance 2 m and within-row distance 1 m). There are 44 patches per plot, *i.e.*, 11 patches per tree species per plot for a total of 1100 patches (Scherer-Lorenzen et al., 2007). The experiment has a gradient of tree functional diversity levels from four similar to four most different tree species. The species pool contained the following 16 species, each with n = 33 to n = 99 patches: *Acer campestre*, n = 55 patches, *Acer platanoides*, n = 99 patches, *Acer pseudoplatanus*, n = 55 patches, *Betula pendula*, n = 88 patches, *Carpinus betulus*, n = 88 patches, *Fagus sylvatica*, n = 33 patches, *Fraxinus excelsior*, n = 77 patches, *Larix decidua*, n = 44 patches, *Pinus sylvestris*, n = 55 patches, *Populus tremula*, n = 55 patches, *Prunus avium*, n = 55 patches, *Quercus petraea*, n = 77 patches, *Sorbus aucuparia*, n = 88 patches, *Sorbus torminalis*, n = 77 patches, *Tilia cordata*, n = 66 patches, and *Ulmus glabra*, n = 88 patches. Based on a cluster analysis using nine above- and belowground functional traits that represent attributes which are indicative for productivity, resource use and nutrient cycling (e.g., leaf type, light requirement, mean annual increment, leaf N concentration, litter C/N ratio), the 16 species were grouped according to functional similarity, and the functional diversity (Petchey and Gaston, 2002) was calculated for all possible 4-species combinations. From those combinations (1,820 total), 24 were selected to represent the gradient of functional diversity (with one species combination planted twice).

### 2.2. Airborne imaging spectroscopy

On 17 June 2021, the Bechstedt and Kaltenborn sites were measured during an airborne campaign within two hours using a Cessna 208B Grand Caravan airplane. In Kaltenborn, four flight lines were combined to one mosaic and in Bechstedt, two flight lines were combined to one mosaic. The average flight altitude was 680 m above ground level, resulting in an average pixel size on the ground of 1 m. The imaging spectroscopy data were acquired using a HyPlant imaging spectrometer. This spectrometer is an airborne instrument consisting of two sensor modules (DUAL and FLUO). We only used the DUAL module that records reflected radiance in 626 quasi-continuous spectral bands in the wavelength range from 374–2504 nm. The spectral resolution is 3.6 nm in the visible (VIS) to near infrared (NIR) and 10.5 nm in the shortwave infrared (SWIR) regionSiegmann et al. (2019). Different processing steps were applied to the raw HyPlant data. First, the data were radiometrically and geometrically corrected using the CaliGeoPro software (Specim Ltd., Finland) to produce georectified at-sensor radianceSiegmann et al. (2019). Measured spectral radiance data were then converted into top-of-canopy reflectance using the ATCOR-4 (Atmospheric and Topographic CORrection algorithm, ReSe Applications GmbH, Switzerland) atmospheric correction approach (Richter and Schläpfer, 2002), as part of a custom-made processing pipeline for data pre-processing (see Siegmann et al. (2019) for a detailed description of HyPlant data and data processing). During the reflectance retrieval, spectra were resampled and certain spectral regions were cut, resulting in reflectance spectra covering the 400-2500 nm wavelength range in 589 spectral bands. In wavelength regions with strong water absorption (e.g., between 1350-1500 nm and 1850-2000 nm), atmospheric transmittance is very low, causing an irradiance and thus reflected radiance signal close to zero. Such low values typically result in very noisy reflectance data. However, ATCOR-4 allows interpolating these absorption bands, by fitting a function considering the shoulder values on the left and right side of the absorption band. This approach results in a smooth reflectance signal. For our data, this interpolation was applied to several spectral regions with larger absorption bands (i.e., for oxygen-caused absorption at 760 nm: bands 221-233 (750-771 nm); for water vapor-caused absorption at 940 nm: bands 306-356 (897-997 nm), at 1100 nm: bands 371-390 (1082-1189 nm), at 1400 nm: bands 415-443 (1331-1488 nm), and at 1900 nm: bands 496-532 (1785-1985 nm)). As these were interpolated values, we excluded these data from the assessment of effects of species identity and diversity. When comparing spectral profiles and using them for classification, we did not exclude them, but reduced the maximum number of bands or PCs to the number of measured bands. If interpolated variables allow for better discrimination, this is a desirable feature for analysis. As a note, we use the word “similarly-spaced” when subsampling the covered wavelength range to indicate that subsampling only considered the band numbers from 1 to 589. That is, distances between subsampled wavelengths were 10/3 times larger in the SWIR than the VIS and NIR regions.

### 2.3. Mapping of the patch level top-of-canopy reflectance data

We stratified our analysis into three levels of aggregation: i) site level (Kaltenborn and Bechstedt, with several plots per site), ii) plot level (*i.e.*, areas of varying species diversity or functional diversity made up of multiple patches), and iii) patch level (*i.e.*, a group of trees belonging to the same species growing as close neighbors). Our smallest level of aggregation was the patches, each with 25 replicate HyPlant pixels representing multiple trees of the same species. The trees were all planted in the same year from the same seed source on flat ground with homogeneous soil conditions across each site(Scherer-Lorenzen et al., 2007). With the spatial resolution of HyPlant (1 m), we could ensure the correct assignment of the 25 replicate pixels to patches as described in the following paragraph.

Patch-specific pixels were extracted based on detailed planting plans and experimental design information. At the Kaltenborn site, we first determined the center and angle of each subplot. To improve geolocation accuracy, airborne data were aligned with high-resolution satellite imagery (*i.e*., GeoBasis-DE/BKG with its data product based on “older–5/7/2020”) available in the Google Earth app, while visual inspection allowed to further refine the subplot center locations and angles. Within each subplot, the central location of each of the 30 (8 x 8 m) patches was then calculated. Given the potential for mixed pixels at patch borders, variability in canopy representation compared to planting design, and potential location discrepancies (on the order of 1 m), we excluded the outer portion of the pixels and extracted only the 25 most central pixels. For each pixel, we assigned the subplot and patch, distances to the center of its patch, coordinates, and the associated reflectance data.

Patch-specific pixels were extracted in a similar way for the Bechstedt site, with modifications to accommodate the specific planting design. The Bechstedt site consists of 25 plots, each with a hexagonal shape (56 x 43.5 m) and containing 44 circular patches (38.5 *m*^2^ area each). Patch locations were determined by combining the Bechstedt planting design, the central location of each plot, and the plot angle. As with the Kaltenborn data, the 25 most central pixels within each patch were extracted for analysis.

### 2.4. Data analyses

#### 2.4.1. Patch-, plot- and species-level spectral variation

For all analyses, we excluded two extreme outliers which corresponded to two patches of *Picea* in plot 9 in the Kaltenborn site (rows 748, 750; Fig. S1). To exclude edge-of-patch effects (which is particularly relevant in cases where the neighboring patch was of a different species), we subset the 25 pixels per patch to the ten most central pixels. Then, to reduce the effects of shadows apparent in the retrieved reflectance data and known to affect data analysis and interpretation (Vögtli et al., 2025), we used the three brightest (sum of reflectance values from 400–2500 nm) pixels of the ten central pixels for every patch (Schneider et al., 2017). We also analyzed data based on all ten central pixels for comparison and found very similar results (Table S1). Including shaded pixels corresponds to using a larger footprint, such as that of a satellite. Furthermore, shaded pixels may relate to architectural differences between species (Vögtli et al., 2025) and thus contribute to species discrimination. Mean reflectance values per patch were used in all subsequent analyses.

To illustrate species-specific spectral reflectance patterns at the Kaltenborn and Bechstedt sites, we then calculated the mean and SD of reflectance across all measured spectral bands for top-of-canopy pixels from each species (Fig. 2).

**Fig. 2.**
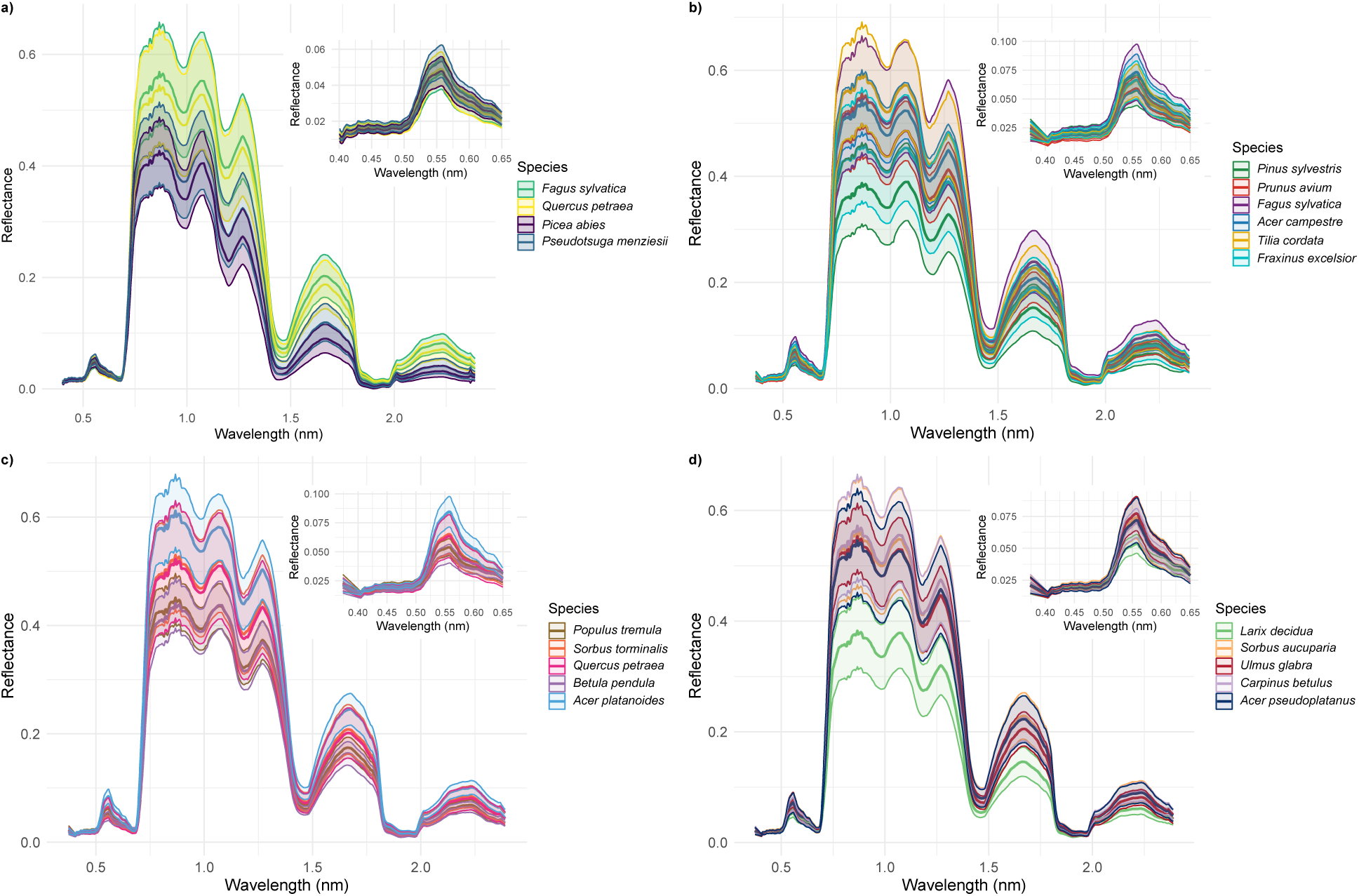
(a) Mean and standard deviation (SD) of the reflectance per species across all experimental plots (n=16) and patches within plots (960 patches in total with 60 patches per plot) at Kaltenborn. Shading indicates one SD of individual reflectance measurements. Inset shows a zoom into the visible range of the spectra. (b–d) Mean reflectance and shaded SD of species at the Bechstedt site with a total of 25 plots and 44 patches within plots; panels b–d are for different subsets of species to facilitate visual distinction of spectra. To visually reflect phylogenetic structure, species are color-coded by taxonomic order with more closely related species sharing similar hues: green tones for Pinales, purple–magenta tones for Fagales, blue hues for Sapindales, red–orange tones for Rosales, brown tones for Malpighiales, yellow for Malvales, and cyan for Lamiales. At both sites, each patch contains multiple trees of a single species. Note that the shown reflectance spectra contain interpolated spectral regions as part of the atmospheric correction.

#### 2.4.2. Analysis of variance (ANOVA)

To determine which spectral wavelengths had the largest discriminatory power between plots, subplots, diversity levels, tree species, and their interactions, we fitted linear models and calculated analysis of variance (ANOVA) tables. Discriminatory power was quantified by contribution to total variance, *i.e.*, percent sum of squares (%SS = increment of multiple R2 when a term is added to the linear model, equivalent to effect size in statistical terminology (Rosenthal and Rosnow, 1985). %SS as an effect size has the advantage that it gives an overall effect size for factors with more than one level (Cohen, 1988). Note that we used the p-values from the corresponding terms in an exploratory rather than confirmatory way and therefore did not correct them for multiple testing. For the Kaltenborn dataset, the model was coded in R as: *y* ~ *logsr* + *DV* + *monoSP* + *PL* + *SPL* + (*CL* + *PI* + *FA*) + *logsr* : (*CL* + *PI* + *F A*)). Plot diversity was coded as a factor with ordered levels 1, 2, 3, and 4 based on the number of species planted (DV), and we also included its log-linear contrast (logSR) as is typically done in the analysis of biodiversity experiments (Schmid et al., 2017). In addition, we used a factorial contrast with four levels that separated the four monoculture plots containing different species from each other (mono SP). Species identity in each patch was coded as a factor (SP=CL+PI+FA) split into classes, distinguishing gymnosperms from angiosperms (CL), and the two species in each class by indicating the presence/absence of *Picea* (PI) or *Fagus* (FA). Plot structure was represented by plot ID (PL) and subplot ID (SPL) within plot ID. Interaction terms were fitted to examine whether differences between classes and species were increased or reduced by the log-linear contrast of diversity. For the Bechstedt dataset, functional diversity for each plot (FD) was characterized by counting the number of functional groups present within the plot based on species composition. Functional group identity was assigned to each species using predefined group labels derived from species-specific trait classifications. This resulted in 1 plot containing a single functional group, 11 plots containing two functional groups, 7 plots containing three functional groups, and 7 plots containing four functional groups. The linear model was coded in R as: *y* ~ *fd* + *FD* + *PL* + (*CL* + *OR* + *SP* )+ *fd* : (*CL* + *OR* + *SP* ). Species were again grouped into the two classes gymnosperms and angiosperms (CL), and for the angiosperms into taxonomic orders within classes (OR), coded as a categorical contrast among the 16 species (SP) to account for above species-, within class-level phylogenetic differences. Note that SP fitted after CL and OR assesses differences among species within orders. The following seven orders were assigned based on established taxonomic classifications. Gymnosperms: *Pinales* (*Pinus sylvestris*, *Larix decidua*), angiosperms: *Malpighiales* (*Populus tremula*), *Rosales* (*Prunus avium*, *Sorbus torminalis*, *S. aucuparia*, *Ulmus glabra*), *Fagales* (*Betula pendula*, *Fagus sylvatica*, *Quercus petraea*, *Carpinus betulus*), *Sapindales* (*Acer pseudoplatanus*, *A. platanoides*, *A. campestre*), *Malvales* (*Tilia cordata*), and *Lamiales* (*Fraxinus excelsior* ). Here, analogous to species richness in Kaltenborn, fd represents a linear contrast of the factor FD. The linear models were fitted using the R function lm() across all 589 wavelengths. We used lm() instead of mixed-model lmer() to obtain %SS values for all terms, but calculated p-values using the mixed-model approach, treating plot ID, subplot ID, and their interactions with taxonomic terms (class, order, species) as random terms. Due to the balanced design of the experiments, results from lm() are the same as those using lmer() (Schmid et al., 2017). The analyses shown in the main text use the three brightest of the ten central pixels per patch.

#### 2.4.3. Classification using linear discriminant analysis (LDA) and a support vector machine (SVM)

We used LDA and a non-linear machine-learning approach (SVM) to assess the discriminatory power of multiple wavelengths combined to identify tree species. We selected an increasing number of wavelengths similarly spread across the reflectance spectrum or an increasing number of principal components (PCs) derived from a principal component analysis (PCA) to fit these LDA and SVM models (up to 439, because 150 spectral bands were removed due to the interpolation during atmospheric correction). Using wavelengths directly maintains the original data space, whereas PCA converts the data space such that the first PCs explain a large amount of the total variation in the data (for Kaltenborn the first 10 PCs explained 99.3% and for Bechstedt 99.2% of the total variation, Fig. S2). Note that PCA does not account for species-level differences in correlations between wavelengths; this potentially leads to a loss of information about differences between species in PC data. LDA then calculates classification functions that are linear combinations of the set of wavelengths or PCs included in the model, while SVM calculates additional variables using kernel radial basis functions from these sets, which allows for non-linear discrimination. LDA models were fitted with the function lda() from the MASS package (Venables and Ripley, 2002) in R (Team, 2026) and SVM models were fitted with svm() function from the e1071 package (Meyer et al., 1999) in R. We used svm() with type = “C-classification” and cost = 10. In the LDA visualizations, we used 68% probability contours because they approximate one standard deviation around each species. This provides a visualization of the central tendency and overlap among species clusters without obscuring patterns through very large contours.

First, we fitted LDA and SVM models using all patches for each of the two experimental sites and with increasing numbers of wavelengths or PCs. Classification accuracy was calculated using confusion matrices with given species identity in columns and predicted species identity in rows.

Second, to assess in-sample prediction accuracy, we used two approaches. For one approach, we randomly shuffled species identities among trees and again calculated prediction accuracies for the shuffled species identities. We found that 20 randomizations were sufficient because standard deviations of accuracies were small. For the other approach, we randomly assigned 120 or 308 patches for Kaltenborn or Bechstedt, respectively, to training sets and the remaining 838 or 792 respective patches to prediction sets and calculated how many trees in these prediction sets were correctly assigned to species. These numbers were used for comparison with the out-of-sample predictions described below. We again found that 20 randomizations yielded small standard errors.

Third, we used patches of particular plots as training sets to predict the species identities of the patches in other plots used as out-of-sample prediction sets. For Kaltenborn, we used plots of a defined diversity level as training sets, with the 120 patches in the two plots with highest diversity, *i.e.*, 4-species mixtures, yielding the best prediction accuracies (and therefore this number was used for the random splits mentioned above). To evaluate whether species classification was influenced by the diversity level represented in the training data, we trained linear discriminant analysis (LDA) models using patches from one diversity level and evaluated their performance on patches from a different diversity level. We generated confusion matrices for all combinations of training and prediction diversity levels to assess how well species could be classified when the diversity context of the training data differed from that of the prediction data (Fig. 7a). The notation *divX → divY* denotes the diversity level of the training data (*divX* ) followed by the diversity level of the prediction data to which the model was applied (*divY* ). Thus, for example, *div1 → div2* indicates that the LDA model was trained using observations from monoculture patches and subsequently used to classify observations from two-species mixture patches. For Bechstedt, we used the 308 patches of plots 1-7 (bottom row of plots in Fig. 1b), containing all 16 tree species, as the training set.

Fourth, we used beech (*Fagus sylvatica*) and oak (*Quercus petraea*), the species occurring at both sites, from patches in Kaltenborn to predict patches in Bechstedt.

For all analyses with training and prediction datasets, we fitted LDA and SVM models only across up to 100 (Kaltenborn) or 200 (Bechstedt) wavelengths or PCs, because prediction accuracy always peaked at smaller numbers. We deliberately used less than half of all tree patches in all training datasets to classify tree patches in prediction datasets, because we were interested in potential applications, where the aim would also be to make large-scale predictions from “ground-truthed” smaller training datasets. All analyses were conducted using patch means of the three brightest pixels and the ten central pixels. However, because differences were small, results using the ten central pixels are only presented in Supporting Information (Table S1). All data analyses were conducted in R (v.4.4.0 and v.4.5.3, R Core Team (2026)).

## 3. Results

### 3.1. Contribution of di!erent wavelengths across the reflectance spectrum to assessing environmental and taxonomic variation

Mean reflectance spectra of the four species in Kaltenborn and the 16 species in Bechstedt using the three brightest pixels per patch are shown in Fig. 2. Species reflectance spectra using the ten central pixels looked very similar (Supporting Information, Fig.S3).

The contributions of species and species-by-diversity interactions to the total variation across the reflectance spectrum for the three brightest pixels per patch are shown in Fig. 3 and Fig. S4S4, respectively. The shown %SS-values represent effect sizes, i.e. amounts of variation in reflectance explained by the corresponding model terms. In Kaltenborn, with a four-species pool, the comparison between angiosperms and gymnosperms was significant across the entire spectrum (indicated by a p-value below 0.05 across all wavelengths), but with an increasing effect size (%SS) with increasing wavelength, particularly beyond the VIS spectral range in the NIR-SWIR spectral region (Fig. 3a). Within the two taxonomic classes, the comparison of *Picea* vs. *Pseudotsuga* was most significant in parts of the VIS that vary with the absorption and reflection of light by non-chlorophyll pigments but with two regions close to peaks of chlorophyll b absorption (centered at ca. 500 nm and 640 nm) being less important; and in the long end of the NIR (750–1200 nm) into the SWIR range (1200–2000 nm). For the differentiation between *Fagus* and *Quercus*, the SWIR above ca. 1400 nm was most significant.

**Fig. 3.**
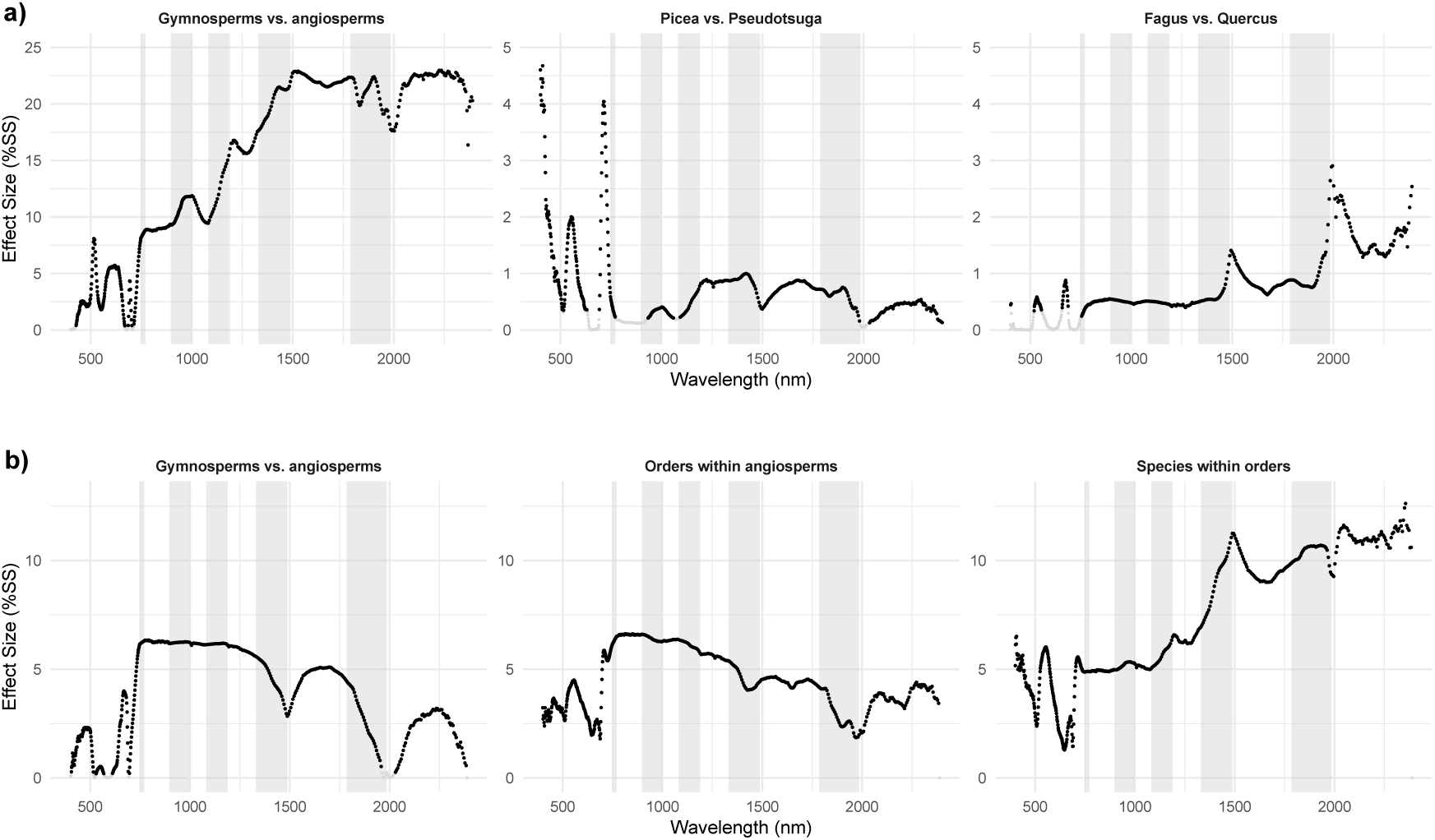
Effect sizes as % sum of squares across the measured spectra for Kaltenborn (a) and Bechstedt (b). The full linear models listed in the statistical analysis section were fitted, but only main effects of species contrasts are shown. Note that the y-axis in (a) is different for the within-class contrasts because these effect sizes were much smaller than those for the between-class contrast. Significant (corresponding P-value < 0.05) effect sizes are shown in black, non-significant effect sizes in gray. Spectral regions interpolated during the atmospheric correction are shown with gray bars in the background. For species-by-diversity interaction terms, which are much smaller, see Fig. S4.

The interactions between taxonomic terms and the logarithm of species richness had effect sizes about an order of magnitude smaller than the main taxonomic differences, but were nevertheless significant for the gymnosperm-vs.-angiosperm contrast above 750 nm (NIR-SWIR, Fig. S4a). In this range, reflectance values between the two classes differed more from each other with increasing species richness. Within the two taxonomic groups, interactions with the logarithm of species richness were only significant between the two gymnosperm species around 500, 1500, and 2000-2500 nm (Fig. S4a). Here, the directionality depended on the specific wavelength region.

In Bechstedt, with a sixteen-species pool and a functional diversity gradient across plots (yet always four species per plot), the differentiation between gymnosperms (*L. decidua*, *P. sylvestris*) and angiosperms was also significant across almost the entire spectrum (Fig. 3b). However, significances in the green to red region of the VIS and at around 2000 nm in the SWIR were lower than in Kaltenborn. The corresponding effect sizes were smallest in the VIS, as for Kaltenborn, but then showed a peak at 750 nm followed by a pattern of declining effect size with increasing wavelength similar to the intensity of a typical vegetation reflectance spectrum (Fig. 3b). Within angiosperms, the six orders were also significantly differentiated across the spectrum, as were the species within orders across gymnosperms and angiosperms. Phylogenetic variation at order level within angiosperms had the largest effect sizes in the NIR, with a pattern similar to the differences between classes. Effect sizes were largest for the comparison of species within orders, especially in the SWIR region, increasing with wavelengths longer than ca. 1200 nm (Fig. 3b).

Also at Bechstedt, taxa responded significantly differently to diversity as shown by the interactions between taxonomic groups and functional diversity, yet again effect sizes were about one order of magnitude smaller than those for the main effects of taxonomic groups (Fig. S4b). Interactions of class with functional diversity were significant in the VIS, especially at blue and red wavelengths in the vicinity of chlorophyll absorption features (Fig. S4b). These interactions indicate that the spectral differences among gymnosperms and angiosperms varied depending on the biodiversity context in which they occurred. As in Kaltenborn, classes were generally better distinguished in plots with higher functional diversity. Interactions of orders with functional diversity were strongest from 750 nm, just outside the VIS, and increased with increasing wavelength throughout the NIR and SWIR. Interactions of the linear contrast of functional diversity with species identity (within orders) were significant throughout the spectrum, with slightly elevated effect sizes in the SWIR between ca. 1400 and 2000 nm (Fig. S4b). These differential responses of taxonomic groups to functional diversity will need further investigation in future studies.

Overall, the ANOVA results indicate that neighboring wavelengths show similar effect sizes, especially in the NIR and SWIR ranges, indicating a dependence that justifies using subsets of similarly-spaced wavelength numbers in the following classification analyses. Similar results were obtained when using all ten central pixels (patch pixels) (Fig. S5). Furthermore, the smaller effect sizes of taxa by diversity interactions suggest that classification should be relatively robust across diversity levels (but see following results).

### 3.2. Distinguishing species using linear and non-linear classification approaches

#### 3.2.1. In-sample classification

Using all data as training data, LDA and SVM models resulted in classification accuracies between 99–100%, except when using wavelengths with SVM (Table 1, Fig. 4, Fig. 5ab). In this case SVM achieved much lower classification accuracies than LDA for both Kaltenborn (89.1%) and Bechstedt (71.5%; Table 1, Fig. 5ab, best prediction for all patches). LDA showed a clear separation of the two classes but also the two species within classes in Kaltenborn (Fig. 4a). In Bechstedt, the Sapindales (three different *Acer* ssp.) were classified closely together and the two conifers overlapped with each other and with *P. tremula* (Fig. 4b).

**Fig. 4.**
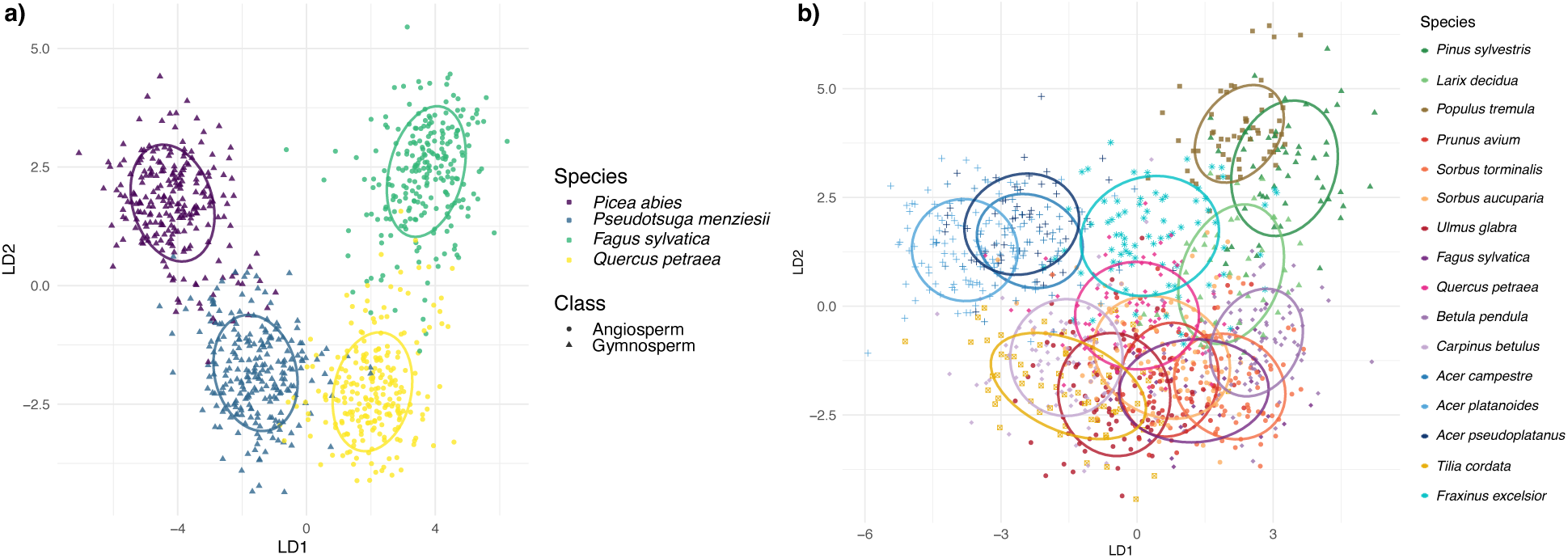
LDA using all tree patches as training data and 439 principal components (PC) derived from reflectance spectra of (a) four species in Kaltenborn (n = 958) and (b) 16 species in Bechstedt (n = 1100). Shown are the first two linear discriminant functions (LD1 and LD2), with patches grouped by species with 68% classification contours. Closely related species share similar hues. In (b) green tones are used for Pinales, purple–magenta hues for Fagales, blue hues for Sapindales, red–orange hues for Rosales, brown hues for Malpighiales, yellow for Malvales, and cyan for Lamiales. In addition, symbol shapes in (b) indicate taxonomic orders: triangles = Pinales, squares = Malpighiales, circles = Rosales, diamonds = Fagales, crosses = Sapindales, squares with a cross = Malvales, and stars = Lamiales.

**Table 1:** Classification performance of linear discriminant analysis (LDA) and support vector machine (SVM) using either similarly-spaced wavelength numbers or principal components (PCs) using the mean of the three brightest pixels. (a) Kaltenborn. (b) Bechstedt. SVM was implemented using the svm() function of the e1071 library in R (cost set to 10; (Meyer et al., 1999)). The standard deviations from multiple simulations (n=20) are shown in brackets. Best predictions per row are shown in bold. The same table for the 10 most-central pixels is shown in the Supporting Information (Table S3).

| Dataset | LDA (wave-lengths) | LDA (PCs) | SVM (wave-lengths) | SVM (PCs) |
| --- | --- | --- | --- | --- |
| **a) Proportion of correct classifications in prediction dataset (958 or 838 patches)** |  |  |  |  |
| 20 wavelengths or PCs |  |  |  |  |
| All 958 patches (same as training dataset) | 0.853 | 0.850 | 0.886 | **0.993** |
| All 958 patches randomly assigned to species | 0.328 (0.015) | 0.325 (0.012) | 0.446 (0.015) | **0.970** (0.004) |
| 120 random training patches, 838 prediction patches (st. dev.) | 0.832 (0.009) | **0.833** (0.009) | 0.764 (0.024) | 0.781 (0.023) |
| 120 four-species mixture training patches, 838 prediction patches | **0.832** | 0.829 | 0.754 | 0.773 |
| 100 wavelengths or PCs |  |  |  |  |
| All 958 patches (same as training dataset) | 0.867 | 0.867 | 0.888 | **1.000** |
| All 958 patches randomly assigned to species | 0.444 (0.014) | 0.447 (0.021) | 0.481 (0.016) | **1.000** (0.000) |
| 120 random training patches, 838 prediction patches (st. dev.) | 0.567 (0.044) | 0.541 (0.060) | **0.773** (0.021) | 0.666 (0.034) |
| 120 four-species mixture training patches, 838 prediction patches | 0.511 | 0.613 | **0.757** | 0.631 |
| Best prediction (wavelengths or PCs) |  |  |  |  |
| All 958 patches (same as classification dataset) | 0.965 (437) | 0.952 (439) | 0.891 (103) | **1.000** (27) |
| 120 four-species mixture training patches, 838 prediction patches | 0.832 (20) | **0.834** (25) | 0.766 (143) | 0.786 (9) |
| **b) Proportion of correct classifications in prediction dataset (1100 or 792 patches)** |  |  |  |  |
| 20 wavelengths or PCs |  |  |  |  |
| All 1100 patches (same as training dataset) | 0.598 | 0.590 | 0.655 | **0.834** |
| 308 random training patches, 792 prediction patches (st. dev.) | **0.494** (0.014) | 0.469 (0.015) | 0.090 (0.009) | 0.437 (0.020) |
| 308 block training patches, 792 prediction patches | **0.453** | 0.404 | 0.278 | 0.336 |
| 100 wavelengths or PCs |  |  |  |  |
| All 1100 patches (same as training dataset) | 0.755 | 0.743 | 0.705 | **0.978** |
| 308 random training patches, 792 prediction patches (st. dev.) | **0.488** (0.015) | 0.453 (0.011) | 0.092 (0.008) | 0.344 (0.012) |
| 308 block training patches, 792 prediction patches | **0.457** | 0.418 | 0.293 | 0.282 |
| Best prediction (wavelengths or PCs) |  |  |  |  |
| All 958 patches (same as prediction dataset) | 0.977 (426) | 0.976 (433) | 0.715 (146) | **1.000** (520) |
| 308 block training patches, 792 prediction patches | **0.489** (49) | 0.448 (50) | 0.306 (402) | 0.356 (35) |

With random reshuffling of species identities for Kaltenborn and using 100 wavelengths or PCs, classification accuracies were around 45% for both wavelength or PCs when using LDA (Table 1). However, using SVM and random reshuffling of species, accuracies were 48% with 100 wavelengths and 100% with 100 PCs. Results were comparable with 20 PCs or wavelengths (Table 1). Because 20 PCs explain 99.6% of variation in multivariate space for Kaltenborn, SVM achieved very high classification accuracy even with the wrong species assignment, as it could separate nearly every individual patch from any other using a sufficient number of non-linear functions in this space. It is likely that with further tuning these accuracies could be improved, but in the case of SVM it seems more reasonable to use PCs instead of wavelengths. Compared with LDA, SVM achieved a high classification accuracy with much fewer PCs, again indicating its ability to distinguish species with non-linear functions within the multivariate space of the first PCs.

Using random splits of the data into training and prediction sets reduced prediction accuracy in Kaltenborn to around 83% with LDA or around 77% with SVM and in Bechstedt to 48.9% or 44.8% when using wavelengths or PCs, respectively, with LDA and to 30.6% and 35.6%, respectively, when using SVM (Table 1, best prediction with training and prediction datasets).

#### 3.2.2. Out-of-sample prediction

The in-sample predictions with random splits of data into training and prediction datasets reported in the previous paragraph had similar classification accuracies as the out-of-sample predictions with patches in the training dataset spatially separate from patches in prediction dataset, both plateauing around 20 wavelengths or PCs (Fig. 5c, d). In the case of Kaltenborn, the best out-of-sample prediction was achieved when we used the two 4-species mixture plots (shown in Fig. 6a), while slightly poorer classifications were achieved with other diversity levels as training sets (Fig. 7a). Our experimental design allowed us to use four different training sets that represented the four levels of species richness in Kaltenborn. Because all species occurred at every species-richness level, each of the training sets contained all species for classification. The main difference was that in the case of monocultures, the four species grew in four separate plots, and in the four-species mixtures they grew together within each of two replicate plots. For the two- and three-species mixtures, the situation was intermediate, that is, two or three species grew together within a plot, whereas the species not occurring in the corresponding mixture grew in a separate plot, yet again in mixtures of two or three species. Similar to the four training sets, we also used these same sets as prediction sets in Fig. 7, so that we could test classification accuracies across diversity levels, *i.e.,* using species growing in monocultures to classify species growing in two-, three-, or four-species mixtures, using species growing in two-species mixtures to classify species growing in monoculture or three- or four-species mixtures and so on. In Bechstedt, out-of-sample classification accuracies were generally low (Fig. 6b).

**Fig. 5.**
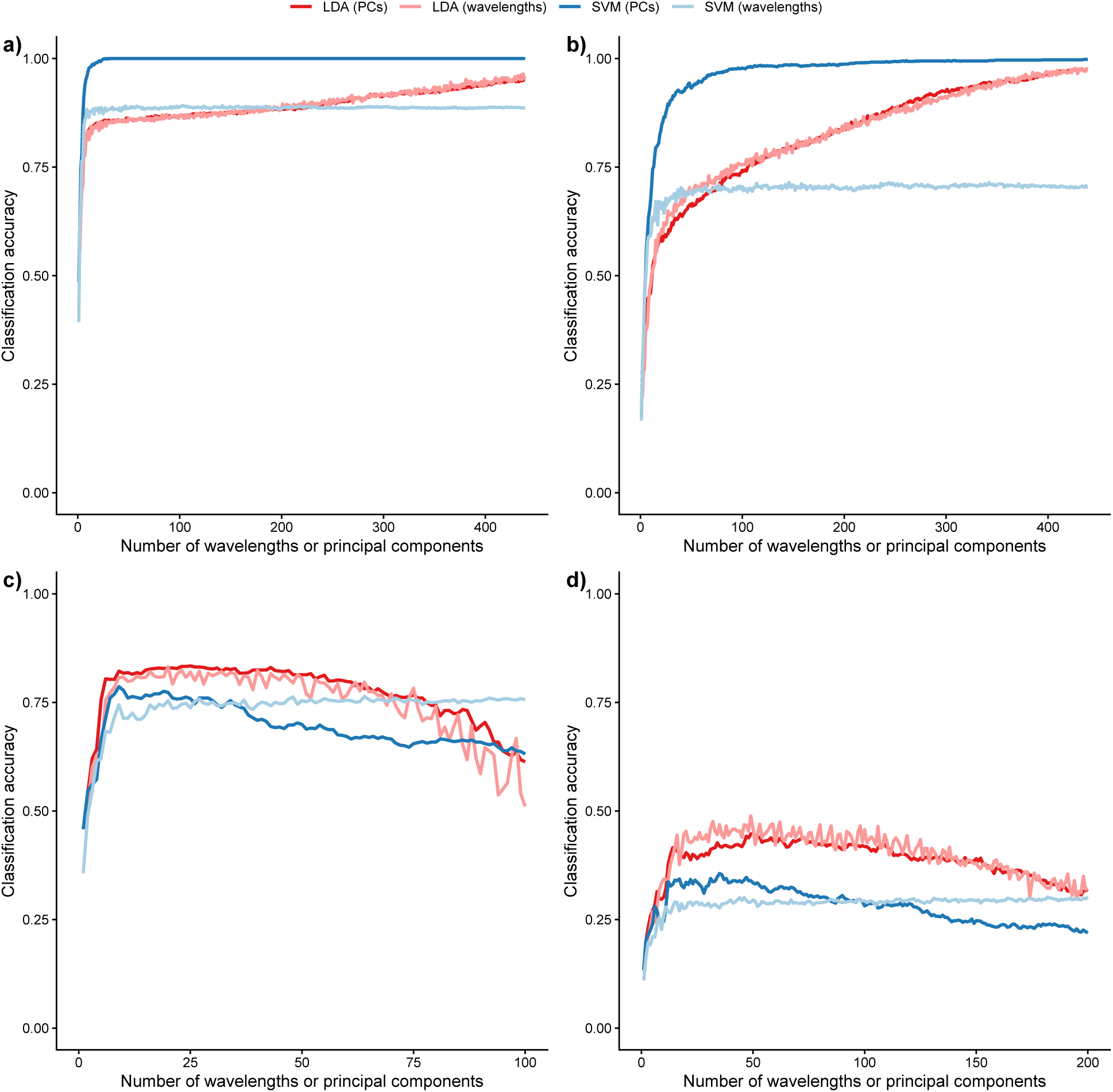
Classification accuracy for four different models, LDA = linear discriminant analysis, SVM = Support Vector Machine. We used 439 similarly-spaced wavelength numbers or principal components (PCs) as input for the models. All models were calculated with the three brightest pixels per patch (see Table S3 for similar results using the 10 central pixels per patch). (a) Kaltenborn, n = 958 patches. (b) Bechstedt, n = 1100 patches. (c) Kaltenborn, out-of-sample prediction, n = 838 patches (training set n= 120 patches). The first 100 PC’s were used, which is why the x-axis is truncated. (d) Bechstedt out-of-sample prediction, n = 792 patches (training set n= 308 patches). The first 200 PC’s were used, which is why the x-axis is truncated. ‘Cost = 10’ in the svm() function.

**Fig. 6.**
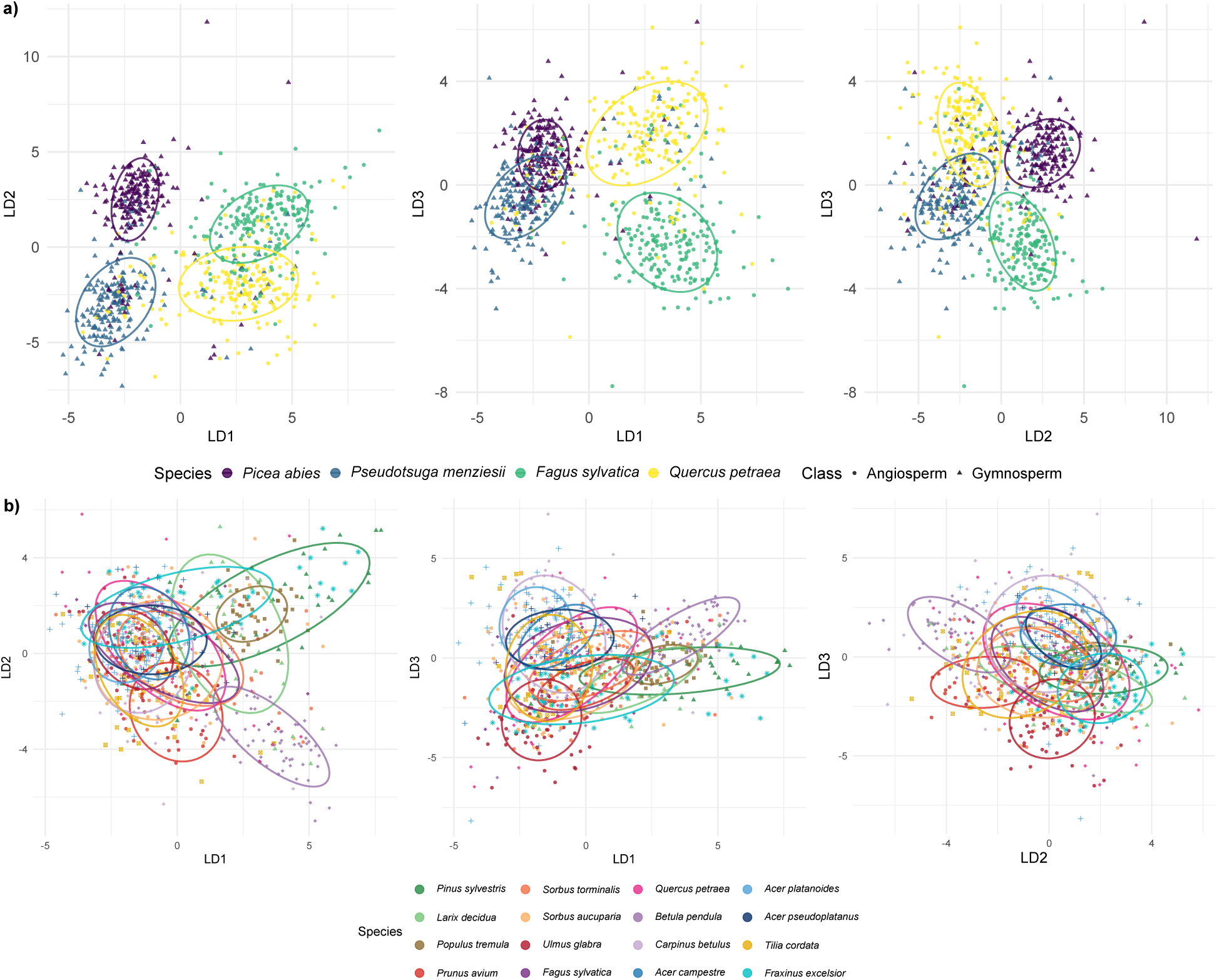
(a) LDA scatter plots for out-of-sample prediction in Kaltenborn, showing species discrimination along LD1 vs. LD2, LD1 vs. LD3, and LD2 vs. LD3 using the optimal number of 25 PCs. The model was calibrated on spectra from plots with diversity level 4 (training set) and applied to plots with diversity levels 1–3 (prediction set). Each point represents an individual monospecific patch of trees from the prediction set. (b) LDA scatter plots for out-of-sample prediction in Bechstedt using the optimal number of 50 PCs. The model was calibrated using data from plots 1–7 and applied to plots 8–16. Each point represents an individual monospecific patch of trees from the prediction set. Closely related species share similar hues. Green hues for Pinales, purple–magenta hues for Fagales, blue hues for Sapindales, red–orange hues for Rosales, brown hues for Malpighiales, yellow for Malvales, and cyan for Lamiales. In addition, symbol shapes indicate taxonomic orders: triangles = Pinales, squares = Malpighiales, circles = Rosales, diamonds = Fagales, crosses = Sapindales, squares with a cross = Malvales, and stars = Lamiales. Ellipses indicate the 68% confidence region for each species.

**Fig. 7.**
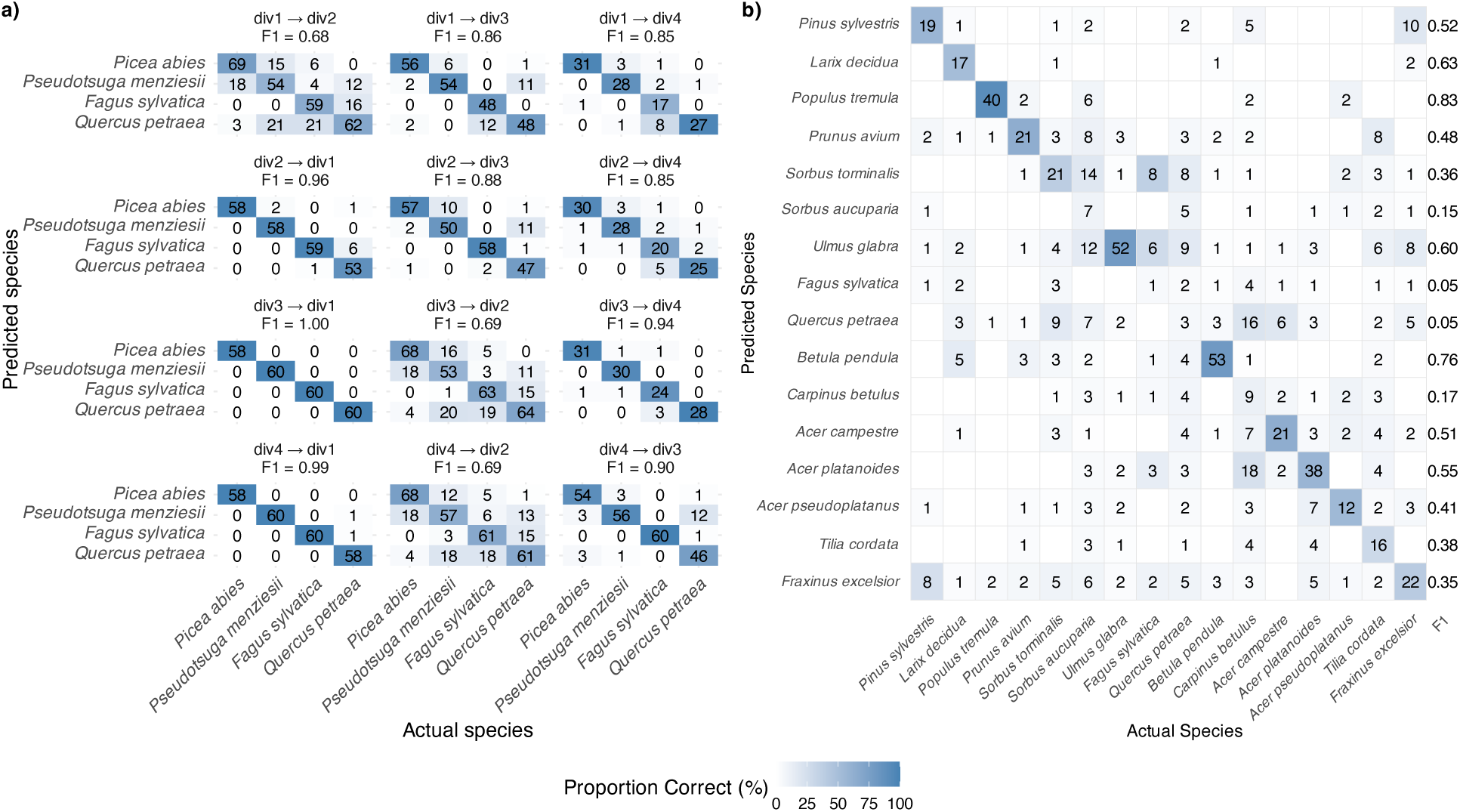
(a) Confusion matrices based on LDA using the first 25 principal components (PC) with different training sets for plots in Kaltenborn; div1 = monocultures, div2 = two-species mixtures, div3 = three-species mixtures, div4 = four-species mixtures. The first term denotes the diversity level of the patches used to train the LDA, and the second denotes the diversity level of the patches used for classification. Thus, div1 → div2, for example, represents an LDA trained on monoculture patches of the four species and predicting those four species in two-species mixtures. Note that the number of replicates per species decreases with increasing species diversity because the total tree density (here patch-density) is held constant. (b) Confusion matrix based on LDA using the first 50 PCs with plots 1–7 as the training set to predict the rest of the plots in Bechstedt. The heatmap indicates the percentage of correctly assigned patches, with darker colors indicating a higher percentage up to 100%. The numbers refer to the actual number of patches. Species are grouped by taxonomic order. F1, the harmonic mean of precision and recall, was calculated as 2 * precision * recall / (precision + recall).

In the case of Bechstedt, we only used one training set (plots 1-7) for prediction. Within the training set, the proportion correct was >95% (Fig. S6). In the prediction set, prediction accuracy was >80% for species such as for example *U. glabra*, *P. tremola*, *B. pendula* and *A. platanoides* (Fig. 7b). However, for some species with lower patch number, the proportion correctly classified was below 25%. Notably, we achieved poor classification of *F. sylvatica*, with no patch being correctly classified (Fig. 7b). Similarly, we could only predict four patches of *Q. petraea* correctly. For *S. aucuparia* only five out of 19 patches were correctly classified.

Using *F. sylvatica* and *Q. petraea* patches in Kaltenborn as training set to predict these species in Bechstedt was less successful than the above out-of-sample prediction within the Kaltenborn experimental site but better than that within the Bechstedt experimental site, yet only a few wavelengths or PCs could be used for discrimination (Table 2).

**Table 2:** Classification performance of LDA and SVM using either similarly-spaced wavelength numbers or principal components (PCs) with the mean of the three brightest pixels (top-of-canopy means) and patches from Kaltenborn as training set to predict *Fagus* and *Quercus* at Bechstedt (prediction set). The wavelength numbers or numbers of PC’s needed for the best prediction are shown in brackets. Best predictions per row are shown in bold.

| Dataset | DA, wavelengths | DA, PCs | SVM, wavelengths | SVM, PCs |
| --- | --- | --- | --- | --- |
| Proportion of correct classifications in prediction dataset (110 patches in Bechstedt) |  |  |  |  |
| 20 wavelengths or principal components, 476 patches in training set from Kaltenborn |  |  |  |  |
| Overall (both species) | 0.664 | **0.673** | 0.609 | 0.645 |
| *Fagus* | 0.303 | **0.424** | 0.000 | 0.364 |
| *Quercus* | **0.818** | 0.779 | 0.792 | 0.766 |
| 100 wavelengths or principal components, 476 patches in training set from Kaltenborn |  |  |  |  |
| Overall (both species) | 0.664 | 0.636 | 0.618 | **0.673** |
| *Fagus* | 0.182 | 0.485 | 0.000 | **0.515** |
| *Quercus* | **0.870** | 0.701 | 0.818 | 0.740 |
| Best prediction (wavelengths or principal components) |  |  |  |  |
| Overall (both species) | **0.782** (393) | 0.746 (206) | 0.700 (10) | 0.736 (182) |
| *Fagus* | 0.940 (4) | **0.970** (3) | 0.697(1) | 0.697 (41) |
| *Quercus* | **0.948** (90) | 0.909 (173) | 0.935 (10) | 0.935 (6) |

For model fitting, SVM performed better than LDA; however, for prediction LDA outperformed SVM (Table 1, Fig. 5). While the use of PCs improved SVM performance (Fig. 5), the original wavelengths remained adequate for LDA (Fig. 5ab). Training accuracy using LDA increased sharply at the beginning but did not level off until almost all wavelengths or PCs were included (Fig. 5ab). Prediction accuracy reached the highest values with fewer wavelengths or PCs (ca. 10-20; Table 1, Fig. 5cd).

The results presented here in the main text refer to the analyses using the three brightest central pixels. Using ten pixels (representing the whole canopy including shaded areas) led to a negligible improvement in classification accuracy using the LDA approach (see Supporting Information, Table S3). For example, in Kaltenborn, the best LDA prediction using the three brightest pixels reached 0.832 (with 20 PCs) and the best LDA prediction using all ten central pixels reached 0.850 (using 20 PCs). For the SVM approach, using 10 pixels vs. 3 pixels did not improve classification and prediction accuracies (Table S3).

## 4. Discussion

Remote sensing has been put forward as a promising technology for large-scale monitoring of forests (Fassnacht et al., 2024) including species, functional and genetic diversity (Schneider et al., 2017; Zheng et al., 2023; Kamoske et al., 2022; Czyż et al., 2020). Substantial progress has been made in mapping biodiversity using both satellite imagery and airborne spectroscopy (Fassnacht et al., 2024). However, challenges related to remote sensing-based tree species classification using high resolution imaging spectroscopy data remain, including the need for sophisticated data preprocessing to account correctly for atmospheric, topographic, BRDF (Vögtli et al., 2024), and shadow effects (Vögtli et al., 2025), and the spectral similarity of different species at canopy level caused by structural interferences partly masking leaf-level information (Torabzadeh et al., 2014, 2019). Further, difficulties exist in disentangling taxonomic and environmental variation in observational studies, where the two likely co-vary (Tian et al., 2024). Here, we used an experiment with planted trees, such that these confounding variables were held constant, to assess the potential to map species richness using airborne spectroscopy under standardized conditions. We show that under these conditions, different tree species show distinct reflectance spectra that can be used for classification in low-diversity settings. However, in more species-rich forests, classification accuracy remains low for out-of-sample prediction of species identities. This confirms existing findings indicating limited tree species classification capability if using optical remote sensing data only (Torabzadeh et al., 2019; Feret and Asner, 2013), and, as we discuss below, this seems to be an inherent information-content limitation that cannot be overcome by more sophisticated classification methods.

### 4.1. Variation in reflectance spectra among taxa and in di!erent diversity settings

Our analysis of variance results showed large taxonomic variation in reflectance spectra and smaller but still significant moderation of this taxonomic variation by species richness in the case of the Kaltenborn site, with a four-species pool, and functional diversity in the case of Bechstedt, with a sixteen-species pool. Thus, variation in the biotic environment influenced species-specific canopy reflectance patterns. This highlights that remote sensing can capture not only taxonomic differences among species but also modifications of species phenotypes associated with biodiversity effects. While classes (gymnosperms vs. angiosperms) and orders (in Bechstedt) were most differentiated, species within these larger taxonomic groups were less differentiated, yet still significantly different throughout the measured spectrum (Fig. 3). That is, gymnosperms and angiosperms differed in structural properties and foliar biochemical composition reported for these groups in the literature. Taxonomic and phylogenetic signals have been observed in previous studies in leaf reflectance spectra (Meireles et al., 2020; Cavender-Bares et al., 2025b) and at canopy scale(Asner and Martin, 2009).

The effect-size spectra in Fig. 3 indicate that adjacent wavelengths in general have similar taxonomic discriminatory power, explaining why similarly-spaced subsamples of wavelengths can achieve accurate classification. The spectral absorption properties of some dominant leaf biochemical constituents (e.g., chlorophyll, water) and the spectral fingerprints of structural canopy properties (e.g., leaf area index, leaf angle distribution) are relatively smooth across wavelengths (Jacquemoud et al., 2009) and can be well captured with a spectral resolution in the nanometer scale as offered by imaging spectrometers (Cavender-Bares et al., 2020), serving a classification scheme as proposed here. It should be noted that imaging spectroscopy also allows to assess more subtle and narrow spectral features caused by specific pigment absorption (e.g., xanthophylls)(Van Wittenberghe et al., 2024) or fluorescence emission (Mohammed et al., 2019), information that could be integrated in forest diversity assessments (cf. (Tagliabue et al., 2020)). The retrieval of such information, however, requires finer spectral resolution, and similarly-spaced subsampling as used here would not be applicable when integrating such information.

In our analysis, we found that variation in taxonomic discriminatory power was larger and on average less significant in the visible range than in the NIR and SWIR ranges (see Fig. 3). Obviously, all species must absorb most radiation in the visible range (see Fig. 2) to maintain photosynthesis, possibly explaining the reduced taxonomic variation in this range. An exception to this was the difference in the chlorophyll absorption range between *Picea* and *Pseudotsuga* at Kaltenborn and the interaction of class-level differences (angiosperms versus gymnosperms) with functional diversity in these spectral regions in Bechstedt.

The observed increase in taxonomic separability toward longer wavelengths is consistent with known spectral sensitivities to leaf internal structure and biochemical composition. Reflectance in the visible range is primarily influenced by pigment absorption, particularly chlorophyll and accessory pigments (Gitelson et al., 2003; Blackburn, 2007), while the near infrared is largely governed by scattering effects associated with internal leaf structure (Gates et al., 1965; Jacquemoud and Baret, 1990). In contrast, the shortwave infrared region contains diagnostic absorption features related to water content and foliar biochemical compounds such as lignin, and cellulose (Kokaly et al., 2009; Asner and Martin, 2008). This region may incorporate signatures of resource use strategies and specialized metabolism and may be less conserved within phyla than are pigments or typical leaf and canopy structural patterns (Meireles et al., 2020) and this may explain the stronger taxonomic differentiation observed in this spectral domain.

As for the responses of classes to species richness at the Kaltenborn site and orders and species at the Bechstedt site these taxonomic groups likely differed in their response to diversity primarily in terms of structural features, water content, and non-pigment constituents such as polyphenolics, proteins, and cellulose. In contract, classes at the Bechstedt site likely differed in their response to functional diversity via changes to their photosynthetic pigmentation. Overall, differences between species in reflectance spectra tended to increase with diversity, suggesting plastic niche separation among species in mixtures, an effect previously described in other forest biodiversity experiments (Niklaus et al., 2017).

### 4.2. Classification accuracy in training datasets

Using linear discriminant analysis (LDA) or a non-linear machine-learning method (SVM), we could accurately classify all individuals, albeit with a large number of spectral dimensions. At Kaltenborn, the SVM achieved perfect classification accuracy when using the first 100 principal components, even when species identities were randomly shuffled (Table 1). This indicates that the classifier learned a unique spectral fingerprint for each individual patch that it could assign to any arbitrary groups (here, randomly shuffled species identities) using a sufficient number of nonlinear functions. With the first 20 principal components, classification accuracy under random species assignment remained very high (97%; Table 1), further supporting the ability of the SVM to separate individual patches in the high-dimensional spectral space using a large number of nonlinear functions. Basically, SVM can thus calculate almost any highly complicated subspace within which given individuals are located. This is not possible with LDA, which can only produce elliptical subspaces. With randomly shuffled species identities, these “artificial” species obviously will locate in large, overlapping elliptical subspaces with correspondingly poor classification accuracy in LDA.

This indicates that using experimental tree plantations in common environments allows nearly perfect or perfect fits even with large samples of around 1000 monospecific patches, about twice the number or spectral wavelengths. High in-sample tree species classification accuracies (80.0% up to 100.0%) have previously been reported in airborne observational studies (Asner and Martin, 2011; Feret and Asner, 2013; Baldeck et al., 2015; Roth et al., 2015; Seeley et al., 2024), yet did not reach the perfection obtained in our experimental study. For example, Seeley et al. (2023) reached 96.0% using SVM but notably in combination with LiDAR data (Seeley et al., 2023). Interestingly, leaf-level (not canopy-level) spectroscopy with LDA in a tropical forest system also resulted in high classification accuracy: 98% using fresh leaves (Hadlich et al., 2025). Leaf-level spectroscopy measurements are generally more controlled contact measurements with a standardized background and light source and thus have a better signal-to-noise ratio and less uncertainty than aerial measurements, though they are lower throughput and cannot capture canopy-level structural features.

Using LDA with similarly-spaced wavelength numbers was as successful or even more successful than using PCs, while using SVM with PCs was clearly more successful than using SVM with the original wavelengths. PCs explain most variation in phenotypic (*i.e.*, spectral) trait space with fewer dimensions than subsampling the original wavelengths, allowing SVM to identify every individual in this PC space, even if the species identities have been completely randomized. However, PCs are calculated under the assumption that all taxonomic groups have the same correlation structure regarding the original traits, *i.e.*, wavelengths, thus blurring some distinctions that might be useful for discrimination (Maaten, 2009). Subsampling original wavelengths from the spectral data avoids this problem and indicates that focusing on high spatial resolution at the expense of lowering spectral resolution may increase discriminatory power in situations where species are mixed at smaller scale than in the present experiment, where they were planted in monospecific patches of 64 *m*^2^ (Kaltenborn) or 38.5 *m*^2^ (Bechstedt).

### 4.3. Classification accuracy in prediction datasets

For the Kaltenborn site with four species in 958 monospecific patches of 64 *m*^2^ each, high classification accuracy could be achieved in prediction datasets even with small training datasets of 120 patches. Using in-sample prediction (random splits of data into spatially mixed training and prediction datasets), LDA yielded an accuracy of 83% with 20 similarly-spaced wavelength numbers or 25 PCs. Classification accuracy was similar for out-of-sample prediction where we used the two 4-species richness plots as training dataset to predict the spatially separate patches of the other richness levels as prediction dataset. This indicates that the monospecific patches in the 4-species richness plots are highly representative and could well predict species identities in plots of lower species richness, in spite of the significant interactions between taxonomic class and species richness found in the analysis of variance (see Fig. S4a). Predicting species identities in the 4-species mixtures using the plots with lower richness-levels as training sets was slightly less successful (Fig. 7a), presumably because here differences between species in the training datasets were confounded with additional differences between plots (which was not the case for the 4-species mixtures as training datasets). That the best predictions are obtained when using the 4-species diversity environment in Kaltenborn supports the notion that a plot with a mixture of species may provide a better reflection of a real world forest as a monoculture plot.

Using SVM resulted in lower out-of-sample classification accuracy of 77% with 143 similarly-spaced wavelength numbers or 79% with 9 PCs. Classification based on linear models may be more generalizable than classification allowing for non-linear relationships, reflecting a trade-off between model generality and precision (Levins, 1968). It is conceivable that with further tuning of SVM, out-of-sample classification accuracy could be improved (and readers are invited to try using the data provided with this paper). However, searching for an SVM that best predicts the concrete out-of-sample set would represent fitting the model to data at a higher level of analysis and depart from the principle that the out-of-sample data should not influence tuning. Similar issues would apply to stepwise LDA (Hadlich et al., 2025; Asner and Martin, 2011), which we also considered but was less successful than LDA using similarly-spaced wavelength numbers or PCs; and also highly inefficient computationally. With stepwise LDA, one could in principle try all possible subsets of original wavelength numbers to maximize classification accuracy, but again at the expense of fitting the model to data.

At the Bechstedt site with 16 species in 1100 monospecific patches of 38.5 *m*^2^ each, out-of-sample classification accuracy with randomly assembled or a spatially aggregated training dataset(s) of 308 patches was below 50%. Using LDA yielded an accuracy of 49% with 49 similarly-spaced wavelength numbers or 45% with 50 PCs. Using SVM again yielded even lower out-of-sample classification accuracy of 31% with 402 similarly-spaced wavelength numbers or 36% with 35 PCs. The lower out-of-sample classification accuracy for the Bechstedt site may be due to the higher number of species, smaller number of individuals per species, smaller patch area, two instead of four overflights, and combinations of these, or further differences between this site and the Kaltenborn site. Compared with the low out-of-sample classification accuracy at the Bechstedt site, identifying the two common species *Fagus sylvatica* and *Quercus petraea* at Bechstedt using the trees at Kaltenborn for training was more successful (78% using LDA with similarly-spaced wavelength numbers, see Table 2). However, identifying *F. sylvatica* was difficult, reflected in the highest accuracy when only 4 similarly-spaced wavelength numbers (94% accuracy) or 3 PCs (97% accuracy) were used in LDA. *Quercus petraea* patches at Bechstedt could quite accurately be predicted with *Q. petraea* at Kaltenborn, also with larger numbers of similarly-spaced wavelength or PCs (see Table 2).

### 4.4. Limitations

We used the three brightest pixels to restrict the analysis to sunlit canopy elements and to avoid misinterpretations caused by shade and related uncertainties in retrieved data products (Damm et al., 2015; Kukenbrink et al., 2019). Selecting only the sunlit pixels is a common and appropriate approach for extracting trait information from aerial imaging spectroscopy data (Schneider et al., 2017; Torabzadeh et al., 2019; Ferreira et al., 2018). However, this strategy reduces within-crown variability and may exclude structural signals that are informative for species discrimination (Vögtli et al., 2025). To account for this, we repeated most analyses using the ten most central pixels per patch, which include both illuminated and shaded pixels. We found similar results derived from the three brightest pixels versus the 10 central pixels of a monospecific tree patch and in few cases even a slightly better performance. This suggests that the thus introduced structural variations are less relevant than other variables such as larger-scale environmental heterogeneity. This could be good news for the use of satellite remote sensing data, where lower spatial resolution implies that such heterogeneity and shadow information are inherent to pixels.

One caveat is of course, that we have an experimental setting that might not be readily transferable to real world conditions. Our data are from a controlled experiment on level ground, while under natural conditions additional structural variation and shading, due for example to greater tree height variation and uneven terrain, can introduce unpredictable effects(Vögtli et al., 2025). Not only for such cases, it is likely that out-of-sample classification accuracy could be improved with additional information from other kinds of remote-sensing, *e.g.* LiDAR data (Torabzadeh et al., 2019; Morsdorf et al., 2020). For example, Torabzadeh et al. (2019) found that combining airborne imaging spectroscopy and laser scanning improved classification results in a temperate forest with a tree species composition overlapping with that of the experiment used here (Torabzadeh et al., 2019). It is important to note that our experimental setting does not use habitat preferences. In real world conditions, species occurrences are often linked to topography, which can aid species classification. Consequently, environmental variables associated with a given tree species may be more similar within species than among species, providing additional information that can aid species classification (personal communication Tiziana Koch). It is therefore conceivable that the high classification accuracies reported for example by (Asner and Martin, 2011; Roth et al., 2015; Seeley et al., 2024) partly reflect the integration of environmental context in addition to the spectral properties of individual trees. Classification studies of trees in real landscapes naturally benefit from environmental context.

The advantage of our study was thus that we could test classification accuracy using remote sensing-based reflectance spectra on a large number of monospecific patches of tree individuals deliberately planted on homogeneous plots, avoiding confounding effects of tree age, overlap between tree crowns of different species, and habitat selection by species potentially resulting in the association of different species with different microhabitats. However, even under our ideal experimental conditions, classification accuracy for closely related species in the case of Bechstedt with the larger species pool was relatively low. This was likely due to limited discriminatory power of canopy spectral properties (reflectance spectra) rather than measurement error or analysis method. Typically, traits used in non remote-sensing classification keys of plant species rely on features such as leaf margin, number of compound leaflets in compound leaves, number of petals or sepals of flowers, seed size and shape, all traits that are hardly reflected (*sic*) in reflectance spectra. Nevertheless, our analysis of variance results do show significant variation in reflectance spectra even at the level of interactions between taxonomic identities and plot diversity levels, suggesting that these signals can be used to assess subtle effects, some even within species.

In conclusion, while reflectance spectra may allow species-level identification of trees in a forest community only under particular conditions and with a low total number of species or low environmental heterogeneity, they can still be used to assess functional diversity (Schneider et al., 2017) under less ideal conditions. Such functional diversity can then be related to forest responses to environmental impacts such as drought events (see Helfenstein et al. (2025)). Reflectance spectra are one tool for forest diversity assessment that should be used in combination with other information from different sources, including other Earth-observation tools as well as field assessments.

## Author contributions

**Sofia J. van Moorsel**: conceptualization, formal analysis, investigation, methodology, visualization, project administration, writing – original draft, writing – review and editing. **Bernhard Schmid**: conceptualization, formal analysis, investigation, methodology, writing - original draft, writing - review and editing **Michael Niederberger**: data curation, methodology, formal analysis, visualization, writing – review and editing **Jeremiah Huggel**: methodology, formal analysis, writing – review and editing **Michael Scherer-Lorenzen**: methodology, writing – review and editing **Uwe Rascher**: methodology, writing – review and editing **Alexander Damm**: conceptualization, methodology, writing – review and editing **Meredith C. Schuman**: conceptualization, funding acquisition, writing – review and editing

## Acknowledgments

This research was funded by the NOMIS foundation grant “Remotely sensing ecological genomics”. We acknowledge funding by the Swiss National Science Foundation (SNSF), Switzerland for the project Fluo4Eco (grant number 197243). The BIOTREE experiment has been established by the Max-Planck-Institute for Biogeochemistry Jena, Germany, and we are very grateful to Prof. Dr. Ernst-Detlef Schulze for making this project possible. The authors thank Dietrich Mackensen and Klaus Hahner (Federal Forestry Office Thüringer Wald—Bundesforstamt Thüringer Wald, Bad Salzungen) for support and site maintenance.

## Data and code availability

Reflectance data and R code will be made publicly available on Dryad and Zenodo upon acceptance of the manuscript.

## Supporting Information

Supporting information can be found online.

## AI statement

During the preparation of this work, the author(s) used ChatGPT (OpenAI) to assist with language editing and improving clarity of the manuscript. The author(s) reviewed and edited the output as needed and take full responsibility for the content of the published article.

