## Supporting Information for "Using a planted tree biodiversity experiment to evaluate imaging spectroscopy for species classification"

*Table of contents.*

**Fig. S1** PCA for Kaltenborn patch means to visualize outliers.

**Fig. S2** PCA for Kaltenborn and Bechstedt using the three brightest pixels per patch.

**Fig. S3** Mean reflectance data using the ten most central pixels per patch.

**Fig. S4** Effect sizes for the interaction terms using the three brightest pixels per patch.

**Fig. S5** Effect sizes based on the ANOVA's using the ten most central pixels per patch.

**Fig. S6** Confusion matrix for the training dataset in Bechstedt using the three brightest pixels per patch.

**Table S3** Classification performance of LDA and SVM using the ten most central pixels per patch.

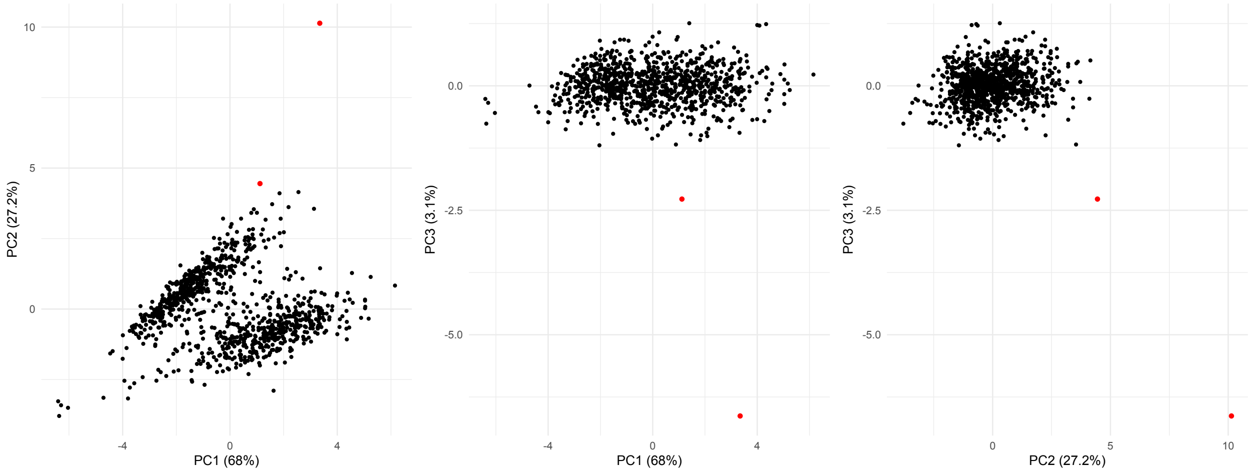


Fig. S1. PCA for Kaltenborn patch means. The removed outliers (*Picea* patches 28 and 30 in subplot m of plot 13) are shown in red.

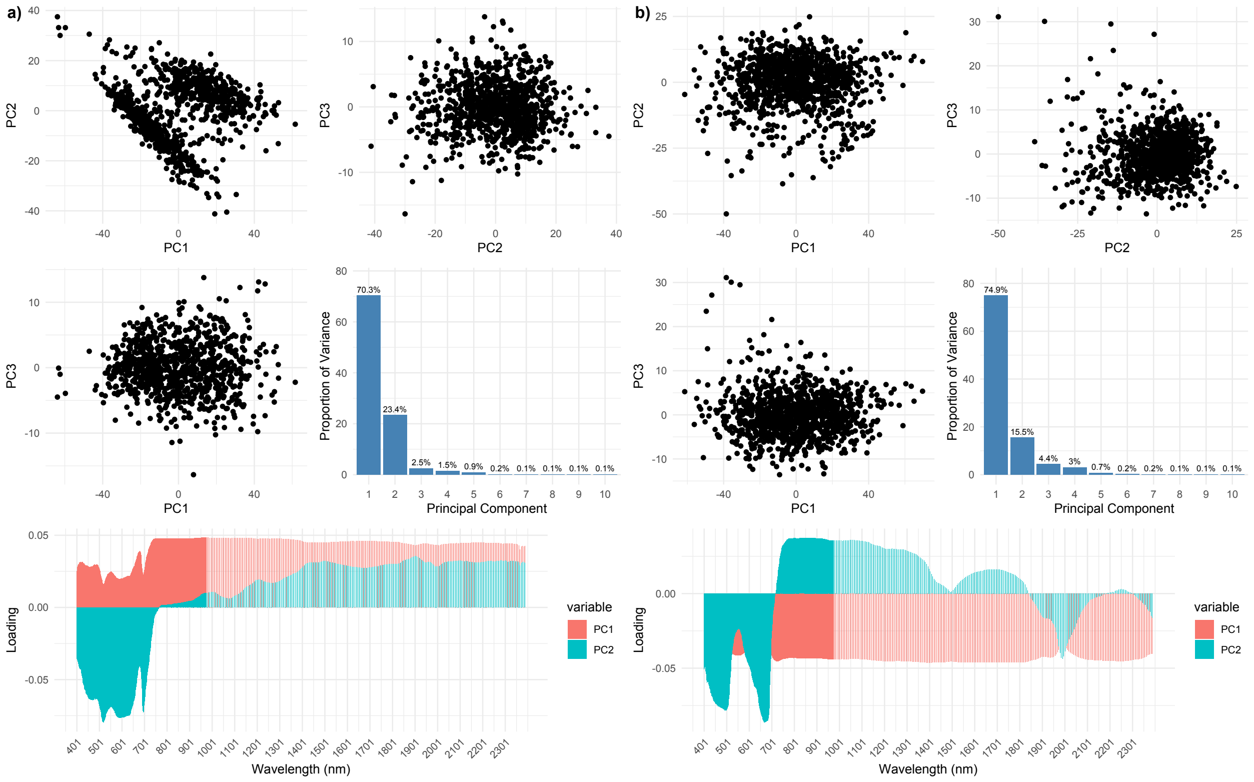


Fig. S2. a) Principal component analysis (PCA) using 439 wavelengths for Kaltenborn. The separation between angiosperms and gymnosperms is visible. Shown is also a scree plot for the 10 first PCs explaining 99.3% of the total variance, with PC1 and PC2 explaining most (93.7%). The percentage explained is indicated above each bar. The bottom panel shows the loadings for the first two PCs across the spectrum. b) PCA using 439 wavelengths for Bechstedt. Shown is also a scree plot for the 10 first PCs explaining 99.2% of the total variance, with PC1 and PC2 contributing most (89.9%). The percentage explained is indicated above each bar. The bottom panel shows the loadings for the first two PC's across the spectrum.

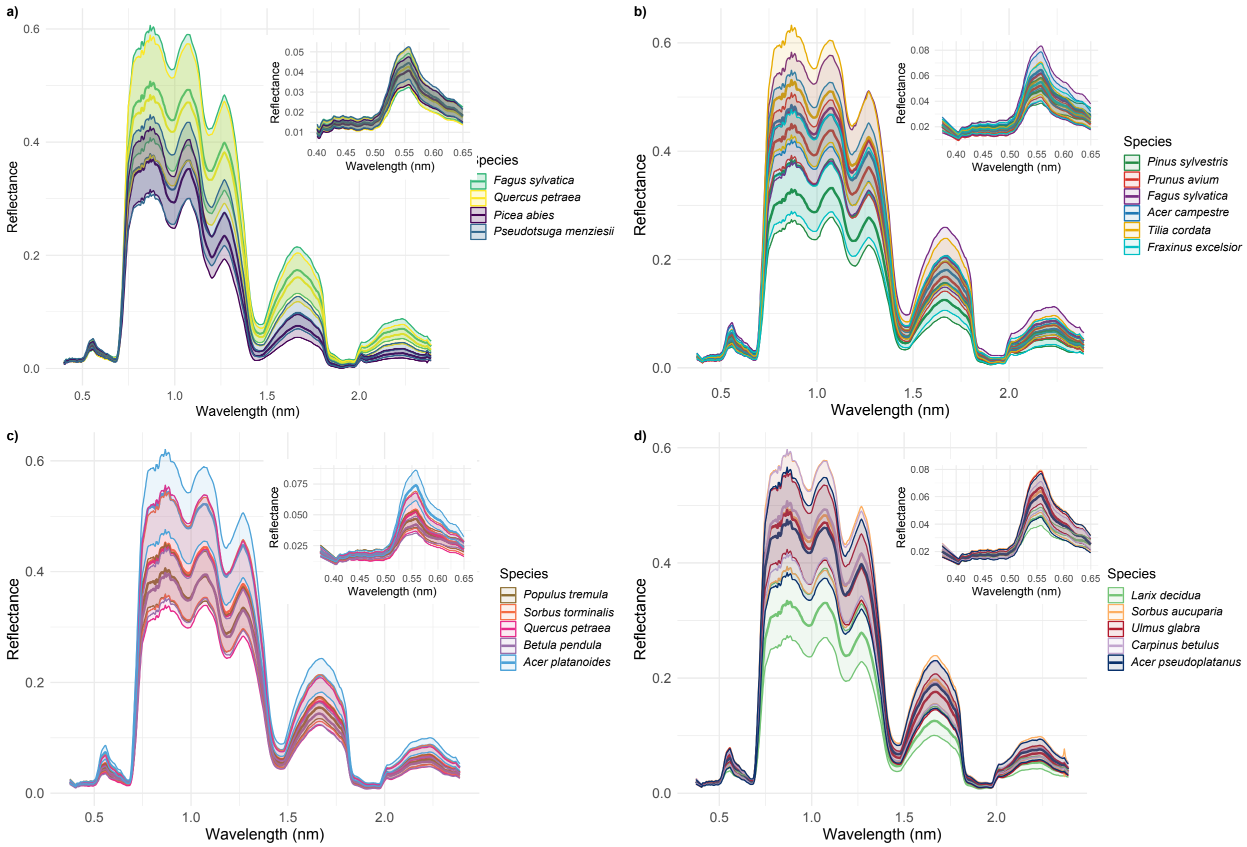


Fig. S3. a) Mean of the reflectance per species and ±1 SD of individual reflectance measurements using the ten most central pixels per patch across all experimental plots (n = 16) and patches within plots (960 patches in total with 60 patches per plot) at Kaltenborn. Inset shows a zoom into the visible range of the spectra. (b–d) Mean reflectance and shaded SE of species at the Bechstedt site with a total of 25 plots and 44 patches within plots; panels b–d are for different subsets of species to facilitate visual distinction of spectra. To visually reflect phylogenetic structure, species are color-coded by taxonomic order. Closely related species share similar hues, while distinct orders are represented by contrasting color groups: green tones for Pinales, purple–magenta tones for Fagales, blue tones for Sapindales, red–orange tones for Rosales, brown tones for Malpighiales, yellow for Malvales, and cyan for Lamiales.

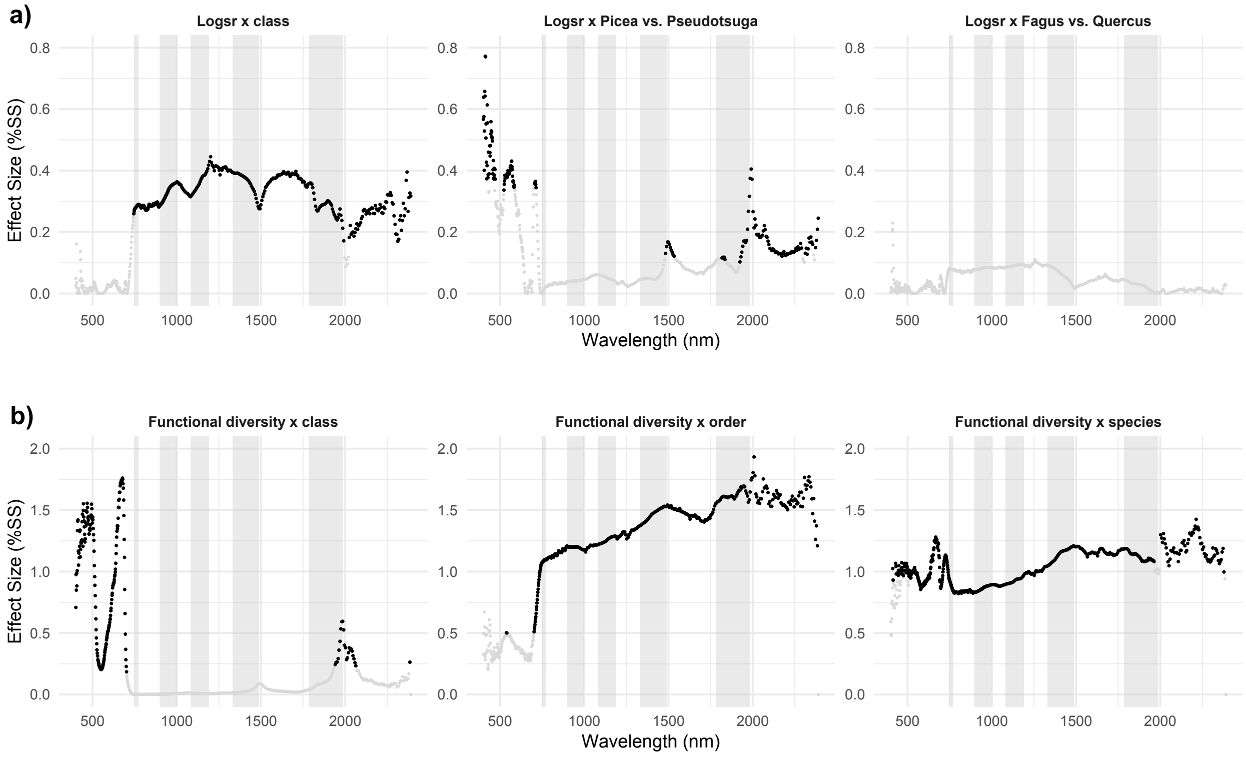


Fig. S4. Effect sizes as percent sum of squares across the measured spectra for Kaltenborn (a) and Bechstedt (b). The full linear models listed in the statistical analysis section were fitted, but here only the interactions of plot diversity (log species richness [Logsr] in Kaltenborn, linear functional diversity in Bechstedt) with species contrasts are shown (main effects of the latter are shown in Fig. 3). Significant (corresponding P-value $< 0.05$) effect sizes are shown in black, non-significant effect sizes in grey. Spectral regions interpolated during the atmospheric correction are shown with grey bars in the background. The three brightest pixels per patch have been used for analysis.

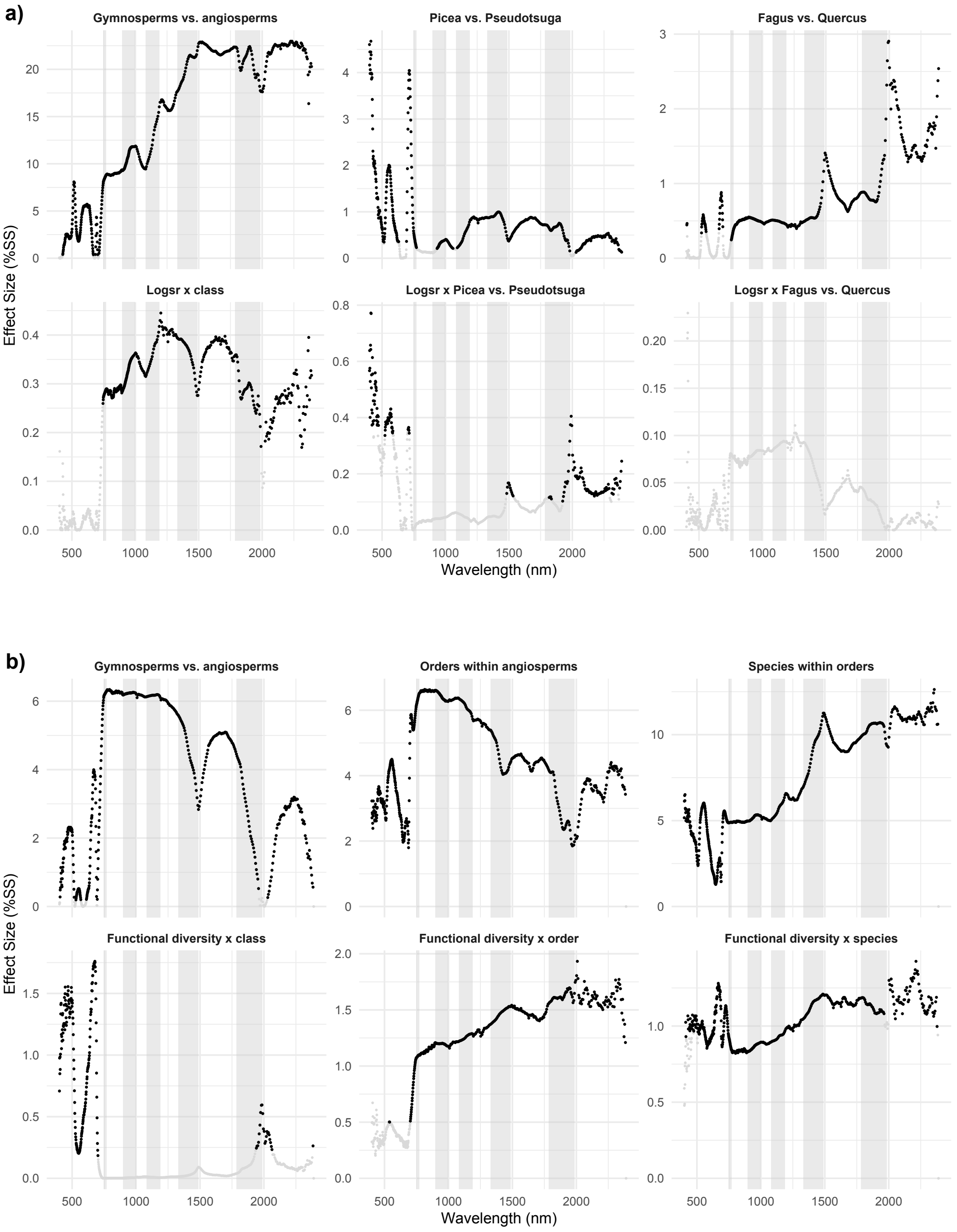


Fig. S5. Effect sizes as percent sum of squares across the measured spectra for Kaltenborn (a) and Bechstedt (b). The full linear models listed in the statistical analysis section were fitted, but only main effects of species contrasts (top row in a and b) and their interactions with diversity (log species richness [Logsr] in bottom row of a and functional diversity in bottom row of b) are shown. Note that the y-axis differs between plots; in particular effect sizes of interactions are an order of magnitude smaller than species main effects. This top rows of this figure correspond to Fig. 3 and the bottom rows to Fig. S4, but here the 10 most central pixels per patch were used for analysis. Significant effect sizes (P-value $< 0.05$) are shown in black, non-significant effect sizes in grey. Interpolated regions are shown with grey bars in the background.

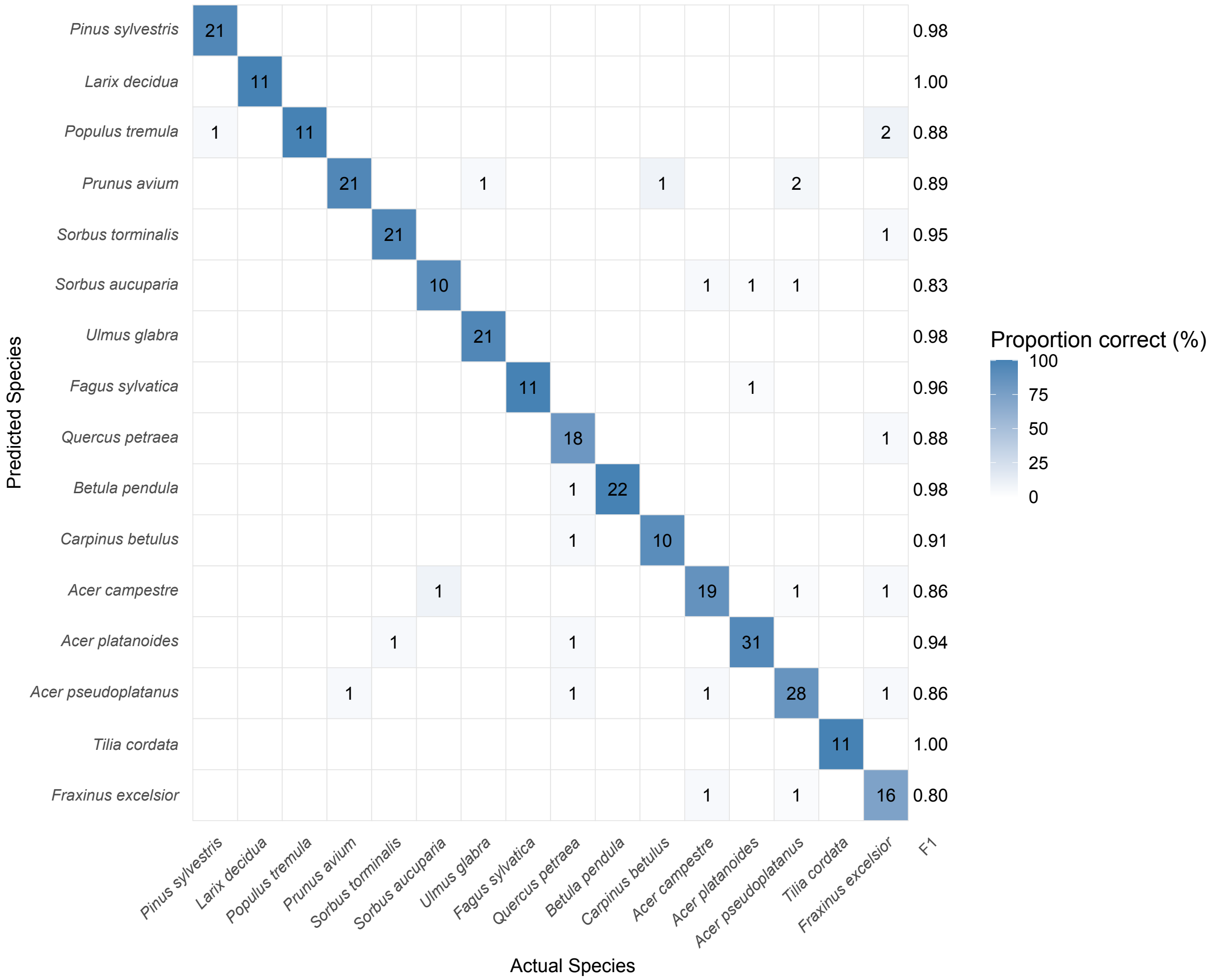


Fig. S6. Confusion matrix based on LDA using the first 50 PCs for the training set in Bechstedt. The heatmap (blue colors) indicates the percentage of correctly assigned patches, with darker colors indicating a higher percentage up to 100%. The numbers refer to the actual number of patches. Species are grouped by taxonomic order. The three brightest pixels per patch have been used for analysis.

Table S1: Classification performance of LDA and SVM using either equally spaced wavelengths or principal components (maximal 439) with the mean of the 10 most central pixels per patch. a) Kaltenborn. b) Bechstedt. LDA: Linear discriminant analysis, SVM: Support Vector Machine

| Dataset | LDA, wavelengths | LDA, PCs | SVM, wavelengths | SVM, PCs |
|---|---|---|---|---|
| a) Proportion of correct classifications in prediction dataset (958 or 838 patches) | | | | |
| 20 wavelengths or PCs | | | | |
| All 958 patches (same as prediction dataset) | 0.857 | 0.863 | 0.843 | 0.915 |
| 120 four-species mixture training patches, 838 prediction patches | 0.850 | 0.833 | 0.712 | 0.760 |
| 100 wavelengths or PCs | | | | |
| All 958 patches (same as prediction dataset) | 0.885 | 0.885 | 0.853 | 0.956 |
| 120 four-species mixture training patches, 838 prediction patches | 0.693 | 0.575 | 0.711 | 0.622 |
| Best prediction (wavelengths or PCs) | | | | |
| All 958 patches (same as prediction dataset) | 0.968 (434) | 0.954 (438) | 0.856 (109) | 1.000 (407) |
| 120 four-species mixture training patches, 838 prediction patches | 0.850 (20) | 0.835 (18) | 0.724 (53) | 0.791 (7) |
| b) Proportion of correct classifications in prediction dataset (1100 or 792 patches) | | | | |
| 20 wavelengths or PCs | | | | |
| All 1100 patches (same as prediction dataset) | 0.642 | 0.616 | 0.440 | 0.834 |
| 308 block training patches, 792 prediction patches | 0.448 | 0.426 | 0.235 | 0.365 |
| 100 wavelengths or PCs | | | | |
| All 1100 patches (same as prediction dataset) | 0.792 | 0.785 | 0.459 | 0.978 |
| 308 block training patches, 792 prediction patches | 0.475 | 0.451 | 0.237 | 0.273 |
| Best prediction (wavelengths or PCs) | | | | |
| All 958 patches (same as prediction dataset) | 0.988 (417) | 0.934 (435) | 0.464 (74) | 0.998 (440) |
| 308 block training patches, 792 prediction patches | 0.538 (58) | 0.491 (57) | 0.241 (25) | 0.396 (19) |
